# *Caenorhabditis elegans* SynMuv B gene activity is down-regulated during a viral infection to enhance RNA interference

**DOI:** 10.1101/2024.07.12.603258

**Authors:** Ashwin Seetharaman, Himani Galagali, Elizabeth Linarte, Mona H.X. Liu, Jennifer D. Cohen, Kashish Chetal, Ruslan Sadreyev, Alex J. Tate, Taiowa A. Montgomery, Gary Ruvkun

## Abstract

Small RNA pathways regulate eukaryotic antiviral defense. Many of the *Caenorhabditis elegans* mutations that were identified based on their enhanced RNAi, the synMuv B genes, also emerged from unrelated genetic screens for increased growth factor signaling. The dozen synMuv B genes encode homologues of the mammalian dREAM complex found in nearly all animals and plants, which includes the *lin-35*/retinoblastoma oncogene. We show that a set of highly induced mRNAs in synMuv B mutants is congruent with mRNAs induced by Orsay RNA virus infection of *C. elegans*. In wild type animals, a combination of a synMuv A mutation and a synMuv B mutation are required for the Muv phenotype of increased growth factor signaling. But we show that Orsay virus infection of a single synMuv A mutant can induce a Muv phenotype, unlike the uninfected single synMuv A mutant. This suggests that decreased synMuv B activity, which activates the antiviral RNAi pathway, is a defense response to viral infection. Small RNA deep sequencing analysis of various dREAM complex mutants uncovers distinct siRNA profiles indicative of such an siRNA response. We conclude that the synMuv B mutants maintain an antiviral readiness state even in the absence of actual infection. The enhanced RNAi and conservation of the dREAM complex mutants suggests new therapeutic avenues to boost antiviral defenses.

## INTRODUCTION

RNA interference (RNAi) is a key eukaryotic defense against viruses and other invading genetic elements. In the nematode *C. elegans*, the RNAi pathway is multifaceted, with an expanded set of 25 Argonaute proteins of many classes compared to the approximately 8 Argonautes from 2 classes present in many animals, including humans. Argonautes bind to small RNAs, including siRNAs or miRNAs of 22 to 26 nucleotides (nt) which guide those Argonaute/siRNA complexes to target mRNAs with complementarity to the siRNAs to induce their degradation or other forms of regulation such as regulation of translation by most miRNAs. In plants, fungi, nematodes, bivalves, and some insects, but not in most animal species, there is a second-stage amplification of siRNAs by the RNA-dependent RNA polymerase (RdRP) proteins. RdRPs use 26nt siRNAs, also known as primary siRNAs, that are generated by the nuclease Dicer to in turn produce a much more abundant set of secondary 22nt-long siRNAs by RdRP primer extensions of primary siRNAs using mRNA templates; these secondary siRNAs often target transposons, integrated viruses, and newly invading viruses. Mutations in a large variety of Argonaute and RdRP genes disable RNAi (1).

Because of this two stage siRNA production pathway, RNAi in *C. elegans* is remarkably potent, and can even be triggered simply by ingestion of double-stranded RNA (dsRNA) produced in an *E. coli* strain engineered using T7 RNA polymerase promoters flanking a cloning site to produce an approximately 1 kb long dsRNA derived from a *C. elegans* genomic or cDNA clone (2). Extensive genetic screens for RNAi-defective (Rde), *C. elegans* mutants revealed loss of function mutations in dozens of genes that disable RNAi, including in the Argonaute protein gene *rde-1* (RNAi defective mutant 1) and a collection of *mutator* genes defined before the discovery of RNAi as mutations that activate transposition of *C. elegans* transposons which are retroviral elements (3, 4). Given that RNAi can be induced even by ingestion of dsRNA, it was somewhat surprising that mutations that enhance the capacity of *C. elegans* to silence mRNAs could be identified: mutations in a distinct set of 20 genes emerged from genetic screens for enhanced responses to dsRNAs that can further enhance the already potent RNAi of *C. elegans* (5–13). A significant fraction of the genes that enhance RNAi (Eri) are members of the synMuv B class genes that were previously identified in totally unrelated genetic screens in the Horvitz and Han labs for increased EGF/Ras signaling in patterning the *C. elegans* cell lineage during development. Among the initial collection of multivulva or Muv mutants from 40 years ago was a set of mutants that carried two loss of function mutations, in *lin-8* <u>and</u> *lin-9*, or *lin-15A* <u>and</u> *lin-15B*, and required loss-of-function alleles of both genes in order to show the Muv phenotype (14–16). Further genetic screens for new mutations that were synthetic Muv with either the *lin-8* (a synMuv A mutant) or *lin-9* (a synMuv B mutant) mutants, but not as single new mutations, revealed a dozen new synMuv B genes that interact with this growth factor signaling pathway. These genes were classified as either a synMuv A (for example, *lin-8* and *lin-56*) or a synMuv B mutation; only a combination of synMuv A plus a synMuv B mutant caused a multivulval phenotype (14).

Molecular analyses of many synMuv B genes mostly by the Horvitz lab showed that many of them encode homologues of the dREAM complex, which is conserved across animals and plants. These include LIN-35, an orthologue of the human tumor suppressor gene retinoblastoma (Rb), and the Rb complex components LIN-53 /RbAp48, LIN-37/MIP40, LIN-54/MIP120, DPL-1/DP, and LIN-9/MIP130 (15–21). Molecular analyses of synMuv A genes showed that *lin-8, lin-56,* and *lin-15A* encode widely expressed nuclear proteins with no obvious homologues outside of nematodes (17, 18); the synMuv A genes either evolve very quickly or have evolved in nematodes or lost in other taxons.

The Muv phenotype of the synMuv A; synMuv B double mutants is caused by abnormally high expression of the growth factor ligand in the hypodermal cells that signal to the adjacent vulval cells. Loss of function mutations in any synMuv B gene in combination with any synMuv A mutant cause excessive LIN-3 EGF growth factor signaling from the hypodermal cells to induce a Muv phenotype in the adjacent vulval cell lineages via the EGF receptor and downstream kinase signaling in those cell lineages (15, 18, 22, 23). Tissue-specific analysis of synMuv A and synMuv B gene activities strongly implicated their function in the hypodermis to regulate the production of the ligand, LIN-3, and for this EGF growth factor signaling pathway in the adjacent vulval cell lineages(24, 25). Tissue-specific expression of the *lin-35* synMuv B gene only in the hyp7 hypodermis is sufficient to rescue the Muv phenotype of a *lin-8; lin-35* double mutant, whereas expression of *lin-35* in the ventral hypodermal cells where the LIN-3 EGF signals are received does not rescue the Muv phenotype (22). In situ mRNA hybridization of *lin-3* shows a dramatic increase in *lin-3* mRNA in the hypodermis only in a synMuv A; synMuv B double mutant, but not in either single mutant (15).

SynMuv B dREAM complex components constitute about 1/3 of the genes that have emerged from screens for enhanced RNAi (5, 9, 10, 12). The previously characterized roles of the synMuv B dREAM complex in cell cycle and transcriptional regulation/chromatin does not obviously intersect with production or response to siRNAs produced during RNAi; the molecular basis of how the DNA-binding proteins and associated proteins of the synMuv B pathway actually intersect with the Argonaute proteins and RdRPs that are central to RNAi has not yet emerged. Further, while the analysis of the synMuv A and synMuv B mutants in EGF signaling from the hypodermis to the ventral precursor cells has been extensive and beautiful, how it connects to the Eri response of the synMuv B mutants is not at all obvious from this molecular genetic analysis of LIN-3 signaling. There are some differences between the hypodermal to vulval cell signaling by *lin-3* and the enhanced RNAi of synMuv B mutants: while both a synMuv A and a synMuv B class mutation are needed to increase LIN-3 signaling from hyp7 for vulval induction, only a synMuv B mutation is required to enhance RNAi (7), and there is no entanglement of the synMuv A mutations in either the enhanced or defective RNAi scenarios.

The pleiotropies of *C. elegans* dREAM complex mutations suggest a function in the intestine and other non-germline tissues on the production of P-granules, the phase-separated granules initially studied by Brangwynne and Hyman, which mediate RNAi, for example Mutator-granules and Z-granules (26, 27). Loss of function mutations in many different *C. elegans* synMuv B genes cause misexpression of these normally germline-specific P-granules in somatic tissues, most dramatically in the polyploid intestine and hypodermis (7, 28, 29). Because RNAi is normally far more active in the germline (30), the somatic activation of the germline P-granule production is consistent with the increased response to dsRNAs that target somatic cells in the synMuv B mutants.

The P-granule misexpression in the somatic intestine is also salient to RNAi because many RNAi components are localized to these perinuclear, phase-separated organelles, and many proteins and RNAs that mediate RNAi localize to P-granules and to the adjacent Mutator granules (31) and Z granules (32), which are also central to RNAi. The gene expression changes in synMuv B mutants supports the model that they regulate particular genomic client loci that mediate RNAi. For example, in L1 stage *lin-35(n745)*/retinoblastoma mutant animals with just two quiescent germline precursor cells (Z2 and Z3) that divide much later in development to generate thousands of germline cells, the P granule genes *pgl-3* (up 28x) and *pgl-1* (up 10x) and the heritable RNAi Argonaute factor *hrde-1* (8x) are significantly upregulated compared to wild type L1 stage (33). These same dramatic gene inductions were also observed in multiple other synMuv B class mutants (28). But this is not a misspecification of intestinal cell fate to germline fate: the animal is viable and the function of the intestine for nutrition and for vitellogenin synthesis and export to the oocyte is intact. This synMuv B mutant intestinal and hypodermal misexpression of P-granules does not depend on a second synMuv A mutation, unlike the increased EGF signaling in the vulval precursor cells (15).

A common theme to the tissues most affected by the synMuv B mutations, the hypodermis and the intestine, is dramatic endoreduplication. The adult *C. elegans* intestine is normally endoreduplicated from diploid at hatching to 32C, 5-inferred full genome duplications; with one duplication at each larval stage but no chromosome condensation or mitosis (25). The hyp7 hypoderm where ectopic P granules are also observed is 8C (34). Thus, two endoreduplicated tissues are affected by dREAM complex genes. Local gene amplification by the dREAM complex has also been observed in Drosophila where the dREAM complex binds at the Drosophila chorion genes to control a 50 kb endoreduplication to nearly 100x the gene dosage (35). This enables the production of the abundant eggshell proteins; mutations in the dREAM complex cause a humpty dumpty phenotype, fragile eggs due to insufficient production of chorion proteins (36–38).

The enhanced RNAi phenotype of the many synMuv B and other Eri mutants, and the siRNA and gene expression changes associated with *eri*-gene mutations points to an elaborate regulation of RNAi activity; the synMuv B mutations which hyperactivate RNAi may place this regulated pathway in a constitutively ON antiviral state, when it is normally a transient state during an actual viral infection. That RNAi may be highly regulated is not surprising: RNAi has evolved by selection for potent antiviral defense during almost a billion years of eukaryotic infection by viruses. But it is surprising that the dREAM complex is so central to RNAi.

Here, we show that *C. elegans* synMuv B mutations activate a viral response pathway, triggering the activation of gene expression and siRNA changes normally activated in a viral infection, but in the absence of an actual viral infection. We show that Orsay RNA virus infection of *C. elegans* causes highly congruent gene expression responses to those of synMuv B mutants. Most importantly, we show that a viral infection induces a physiological state that down-regulates synMuv B gene activity: like the synthetic multivulval phenotype that is normally only observed when *C. elegans* carries both a synMuv A and synMuv B mutation, Orsay viral infection of a synMuv A mutant causes a partially penetrant Muv phenotype. This suggests that the downregulation of synMuv B gene activity or dREAM function is a key defense response that is triggered during an actual viral infection. Thus, the mammalian-conserved synMuv B pathway highlights new avenues to activate production or response to siRNA and may be targeted with drugs to activate human antiviral responses or responses to RNAi-based pharmaceuticals.

## RESULTS

### synMuv B null mutants turn on many genes that are up-regulated during viral infection

Enhanced RNAi (Eri) mutants were first identified by their activation of RNAi in neurons, for example, which normally do not recapitulate the phenotypes caused by loss of function mutations, for example in synaptic components with strong movement disorders (39). The *eri* mutants also show enhanced responses to a standard set of feeding dsRNAs (*lin-1, hmr-1, lir-1, cel-1, unc-73, dpy-13*), that do not cause phenotypes in wild type but strong phenotypes in *eri* mutants (10). One model for the existence of the *eri* mutants is that each of them constitutively activates what is normally a highly transient state of increased RNAi during a viral infection. We explored whether the Eri response of synMuv B mutants might be evidence of their involvement in the orchestration of a coordinated genetic response towards a perceived viral threat, rather than a mere byproduct of gene expression-related abnormalities.

To explore this hypothesis, we analyzed mRNA seq data sets for *C. elegans* synMuv B *lin-35(n745)* and *lin-15B(n744)* null mutants (available through the NCBI GEO database collection), for gene expression signatures that may show similarities to the gene expression responses to infection of *C. elegans* by the Orsay RNA virus. We analyzed the mRNA seq data sets of developmental-stage matched wild-type N2 and synMuv B *lin-35* and *lin-15B* mutants for the induction of genes that are strongly induced during Orsay viral infection (GEO accession numbers in Materials and Methods). About 80 *C. elegans* genes are 3 to 10-fold up-regulated when animals are infected by the Orsay RNA virus compared to control animals (40). A highly overlapping set of 50 genes are 4 to 10-fold up-regulated in *lin-15B* or *lin-35* L1 stage animals compared to wild type L1 animals. For example, 20 of the 50 *lin-15B* mutant induced genes and 22 of the 50 *lin-35* mutant induced genes are also induced in wild type animals upon Orsay virus infection. Many of the most highly virus-induced *C. elegans* response genes and most highly induced *lin-15B* or *lin-35* mutant response genes are the *pals* genes (41, 42), which are also dramatically upregulated in both *lin-35(n745)* and *lin-15B(n744)* null mutants [See **Table 1 and S1 Fig**]. Each of the induced genes shown in Table 1 and S1 Fig are 5-fold or more induced in the synMuv B mutants compared to wild type controls, in the absence of any viral infection. We list the top 50 genes that are up by at least 5-fold in *lin-15B(n744)* and *lin-35(n745),* mutants in **S1 – S3 tables**. The 39 PALS genes are localized in five, 20 to 50 kb clusters within the *C. elegans* genome (41–44). Significantly, a mutation in the *pals-22* gene causes increased transgene silencing, a common response to many enhanced RNAi mutants, including the synMuv B mutants (41–44). The clustering of these genes may be related to their potential co-regulation or co-amplification during a response to a viral infection.

**Table 1.** Differentially expressed *C. elegans* genes upon Orsay-virus infection are upregulated in *lin-35(n745)* and *lin-15B(n744)* synMuv B mutants under non-infection conditions. Shown here are how the top 17 *C. elegans* genes induced after Orsay-virus infection and their level of expression within *lin-35(n745)* or *lin-15b(n744)* synMuv B mutants under non-infection conditions within the mRNA seq data sets *GSM4697089-lin-35[JA1507(n745) rep1/L1, GSM4697090-lin-35[JA1507(n745) rep1/L1], GSM1534086-lin-35[JA1507(n745) rep1 L3 and GSM1534087-lin-35[JA1507(n745) rep1 L3] and GSM4697102-lin-15b(n744)-rep1/L1, GSM4697105-lin-15b(n744)-rep2/L1*], that were obtained from the NCBI Geo collection. The log2 gene expression Fold changes shown for N2 animals upon Orsay virus infection were obtained from Chen et al., 2017 (see ref 40).

| Gene | log2 FC in<br>lin-35(n745),<br>L1 stage | FDR | log2 FC in<br>lin-35(n745),<br>L3 stage | FDR | log2 FC in<br>lin-15b(n744)<br>L1 stage | FDR | Log2 FC<br>in WT(N2),<br>upon Orsay<br>virus infection | FDR |
| --- | --- | --- | --- | --- | --- | --- | --- | --- |
| **B0507.10** | 4.52 | 1.51E-05 | 5.87 | 6.90E-04 | 4.46 | 1.07E-04 | 5.80 | 1.41E-17 |
| **B0507.8** | 4.93 | 3.69E-05 | 6.57 | 1.15E-03 | 4.22 | 8.69E-04 | 6.45 | 4.01E-31 |
| **pals-6** | 3.24 | 1.37E-04 | 5.41 | 6.75E-04 | 3.04 | 1.78E-03 | 4.09 | 9.08E-50 |
| **F26F2.4** | 3.08 | 1.17E-02 | 9.47 | 5.38E-03 | N/A | N/A | 7.79 | 4.09E-116 |
| **sdz-6** | 6.12 | 5.14E-03 | 8.72 | 1.27E-02 | 5.26 | 5.82E-02 | 5.63 | 2.52E-47 |
| **F26F2.2** | N/A | N/A | N/A | N/A | N/A | N/A | N/A | N/A |
| **F26F2.3** | N/A | N/A | 7.13 | 2.69E-05 | N/A | N/A | 8.37 | 6.06E-127 |
| **C43D7.4** | 4.58 | 4.57E-03 | 7.96 | 1.89E-02 | 3.42 | 1.20E-01 | 9.99 | 1.22E-61 |
| **pals-28** | 3.98 | 4.44E-03 | 6.60 | 1.84E-02 | 4.99 | 4.52E-03 | 8.31 | 8.32E-55 |
| **F26F2.5** | N/A | N/A | 8.59 | 2.10E-03 | N/A | N/A | 7.45 | 1.64E-58 |
| **F42C5.3** | 4.80 | 1.33E-05 | 8.59 | 2.10E-03 | 4.37 | 2.00E-04 | 6.49 | 9.12E-90 |
| **pals-5** | 7.59 | 8.19E-07 | 7.89 | 3.47E-04 | 9.06 | 5.80E-07 | 6.72 | 2.47E-65 |
| **F26F2.1** | N/A | N/A | 4.87 | 5.55E-04 | N/A | N/A | 5.76 | 2.52E-62 |
| **eol-1** | 4.48 | 8.42E-07 | 4.67 | 2.45E-04 | 3.89 | 1.80E-05 | 6.06 | 2.32E-79 |
| **Y57B8A.39** | N/A | N/A | N/A | N/A | N/A | N/A | N/A | N/A |
| **C49C8.2** | N/A | N/A | N/A | N/A | N/A | N/A | N/A | N/A |
| **C43D7.7** | N/A | N/A | N/A | N/A | N/A | N/A | 8.85 | 1.39E-32 |

A *pals-5* promoter fusion to GFP has been developed as a reporter of viral infection response by the Wang and Troemel labs after their RNA seq of viral and microsporidia infection showed that *pals-*genes are highly induced (41). The strong induction of *pals-*genes in the *lin-15B* or *lin-35* synMuv B mutants may report the aberrant induction of antiviral defense in the absence of a real viral infection in these mutants. For example, a *pals-5p::GFP* fusion gene is not expressed in wild type *C. elegans* not infected with Orsay virus, but is strongly activated in the intestine of a wild type animal carrying the *pals-5p::GFP* after Orsay virus ingestion, the mode of *C. elegans* Orsay viral infection (41). To explore this viral response in the synMuv B mutants, we used RNAi or mutations to disrupt particular synMuv B genes in a strain carrying *pals-5p::GFP*. RNAi of the synMuv B genes *lin-9* or *lin-13* strongly activated *pals-5p::GFP* [**Fig 1 A&B**]. We also crossed strong loss-of-function mutations of *lin-35(n745)* or *lin-52(n771)* or *lin-9(n112*), into a strain carrying *pals-5p::GFP* and found that all the three synMuv B homozygous mutants robustly activated *pals-5p::GFP* [**Fig 1C**]. We further tested by RNAi the synMuv A genes *lin-8* or *lin-56* as well as several *lido* (<u>LI</u>N-8 <u>do</u>main containing) genes, paralogues of the synMuv A gene *lin-8* (45), but these gene inactivations of synMuv A genes or their paralogues did not induce *pal-5p::GFP* [**Fig 1B**]. These findings support the hypothesis that the *C. elegans* synMuv B mutants may place the animal in a physiological state that is normally induced by a viral infection, triggering the activation of their antiviral response, for example induction of specific viral defense genes and activation of RNAi.

**Fig 1.**
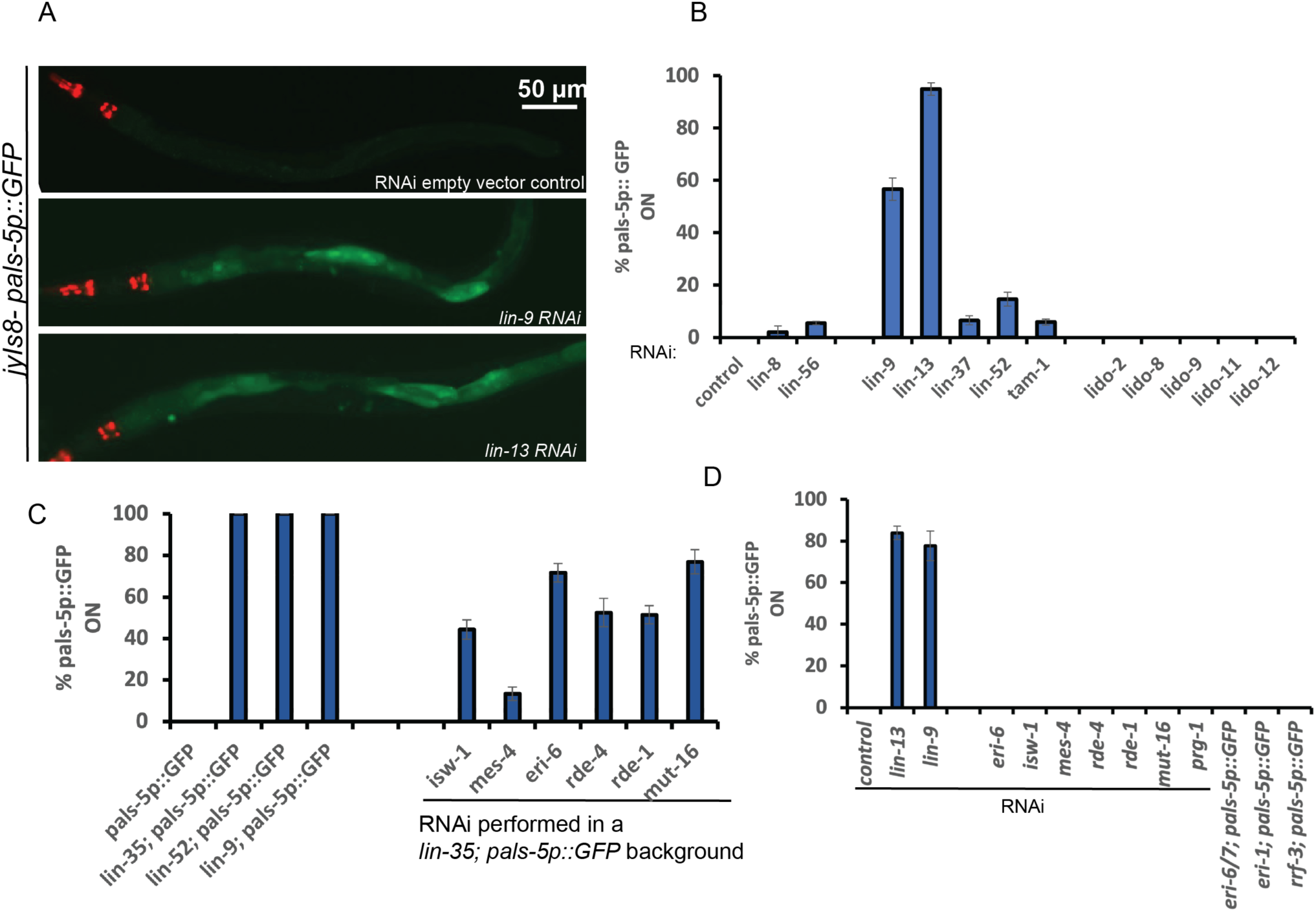
A *pals-5p::GFP* fusion gene is induced in synMuv B Eri mutants even in the absence of an actual viral infection. (A). Fluorescent micrographs showing *jyIs8[pals-5p::GFP],* expression in animals raised on *E. coli* expressing dsRNA of either the L4440 (empty vector control), or dsRNA targeting *lin-9* and *lin-13* synMuv B genes. Scale bar is indicated. (B). Quantification of the percentage of animals showing *pals-5p::GFP* expression when raised on *E. coli* expressing dsRNA against a panel of different synMuv B *eri-*genes including *lin-9*, *lin-13* and non-Eri synMuv A genes *lin-8*, *lin-56,* and a panel of *lido (for* **li**n-8 **do**main) genes (C). Quantification of the percentage of animals showing *pals-5p::GFP* expression in *lin-35(n745);pals-5p::GFP, lin-9(n112);pals-5p::GFP* and *lin-52(n771); pals-5p::GFP* synMuv B mutants and percentage of *lin-35(n745);pals-5p::GFP* animals showing *pals-5p::GFP* expression when raised on *E. coli* expressing dsRNA against the synMuv Eri suppressor genes *isw-1* and *mes-4* and other key genes that encode components central to RNAi such as *rde-1*, *rde-4 and mut-16* (D). Percentage of animals showing *pals-5p::GFP* expression when raised on *E. coli* expressing dsRNA against the underlined panel of genes in the *jyIs8[pals-5p::GFP] animals.* Non-underlined genes are genetic doubles that were generated in the *pals-5p::GFP* background with strong loss-of-function mutations of the genes shown.

Consistent with the *pals* gene-family involvement in anti-viral defense, multiple *pals-22* null mutations were identified in a genetic screen for silencing of multicopy transgenes, which are recognized by the RNAi pathway as foreign genetic elements akin to viruses and silenced by the pathway (44). Transgenes are recognized as foreign, most especially in Eri mutants such as the synMuv B and other Eri mutants (10). The *pals-22* allele that emerged from a screen for transgene silencing also showed enhanced RNAi with the *dpy-13* and *cel-1* feeding RNAi that are used to test for enhanced RNAi (44); thus, the *pals-22* loss of function mutant is an Eri mutant. And the induction of the *pals* genes by viral infection and by the synMuv B mutations in *lin-15B* or *lin-35*, and the observed enhanced RNAi of *lin-15B* or *lin-35* null mutants is consistent with association between *pals-*genes and antiviral defense.

### Suppressors of the synMuv B; synMuv A double mutant Muv phenotypes also suppress *pals-5p::GFP* induction by synMuv B loss of function mutations

The Han group discovered 32 gene inactivations that can suppress the Multivulvae (Muv) phenotype of synMuv A; synMuv B double mutants (14). Seven of those Muv suppressor genes, *mes-4, isw-1, gfl-1, mrg-1, zk1127.3, m03c11.3, c3410.8* were complete suppressors of the Muv phenotype by RNAi. A subset of these synMuv suppressors, also suppress the Eri phenotype of synMuv B mutants (14). Two of the more potent synMuv B Eri suppressors include *isw-1,* which encodes a member of the ISW1 complex and *mes-4*, which encodes a Set domain containing protein (14). We tested whether the induction of *pals-5p::GFP* in synMuv B mutants is functionally linked to their Eri phenotype. We reasoned that suppressing the Muv or the Eri phenotype of synMuv B mutants could also cause a decrease in the induction of the *pals-5p::GFP* reporter. To test this, we targeted *isw-1* and *mes-4* Muv suppressors by RNAi in the *lin-35(n745); pals-5p::GFP* strain. As controls, we also targeted other core components of the RNAi machinery such as the Argonaute gene *rde-1,* the RNA helicase gene *rde-4* and the mutator protein gene *mut-16* and determined the percentage of GFP-positive animals in their F1 progeny. We found that knocking down *isw-1*, *mes-4, rde-1*, *rde-4 and mut-16* by RNAi partially suppressed the percentage of GFP-positive animals in the *lin-35(n745); pals-5p::GFP* animals [**Fig 1C**]. We also determined that inactivation of *isw-1, mes-4, rde-1, rde-4 or mut-16* by RNAi in an otherwise wild type background does not activate *pals-5p::GFP* [**Fig 1D**].

We also characterized the induction of *pals-5p::GFP* in enhanced RNAi mutants that are not synMuv B genes and found that only synMuv B mutations or synMuv B gene RNAi induce *pals-5p::GFP*. For example, inactivation by RNAi or by mutations of the *eri* genes *eri-6, eri-6/7, eri-1,* and *rrf-3* does not activate *pals-5::GFP* [See **Fig 1C & D**]. This suggests that the Eri phenotype induced by synMuv B gene mutations may enhance RNAi via a pathway that is distinct from the other known Eri mutants. In addition, targeting *prg-1*, the worm ortholog of PIWI, which silences integrated viruses in Drosophila (46), by RNAi, also did not activate *pals-5::GFP* expression [See **Fig 1D**], suggesting that piRNA mediated defense response is distinct from that of the Eri phenotype caused by synMuv B mutants.

We also infected *pals-5p::GFP***-**wild type and *lin-35(n745); pals-5p::GFP* animals with the Orsay virus and examined their F2 progeny for *pals-5p::GFP* activation. We examined F2 progeny to allow sufficient time for the viral infection to become well-established within the population. The *pals-5p::GFP* animals not infected with Orsay virus did not induce GFP expression, whereas *pals-5p::GFP* animals infected with the Orsay virus showed a robust activation of *pals-5p::GFP* [**Fig 2A &C**]. The *lin-35(n745); pals-5p::GFP* animals show 100% penetrant GFP activation under both infected and uninfected conditions [**Fig 2A, B & C**]. Inactivation of *isw-1* and *mes-4* by RNAi in *lin-35(n745); pals-5p::GFP* animals under a non-infected state significantly suppressed the *pals-5p::GFP* reporter activation [**Fig 2A, B and D**]. However, upon infection by the Orsay virus, we no longer observed this suppression, which suggests that *isw-1* and *mes-4* function while required for the induction of *pals-5p::GFP* in *lin-35(n745)* null mutants under non-infection conditions may be dispensable or insufficient in the context of an actual viral infection.

**Fig 2.**
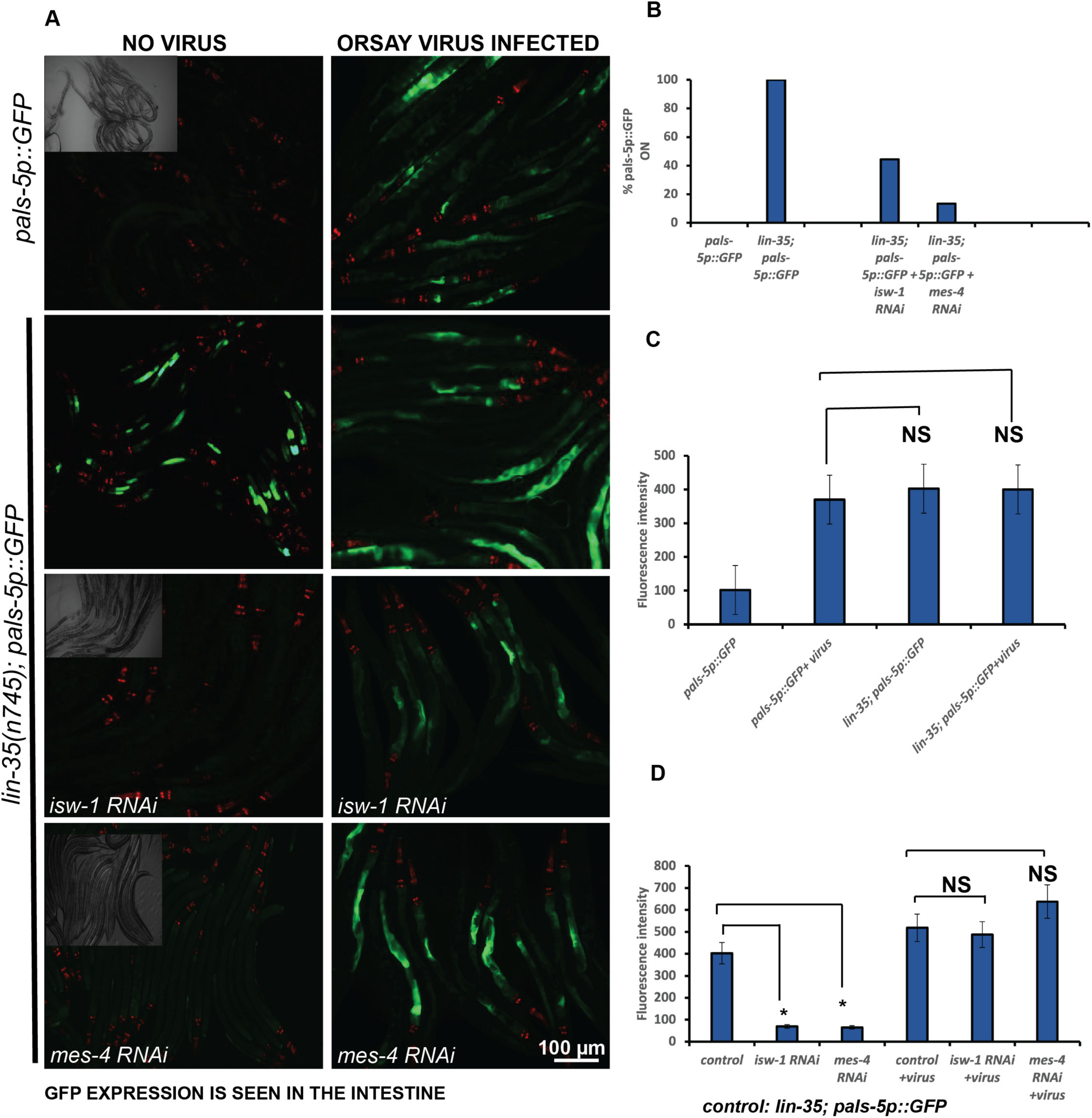
Gene inactivation by RNAi of the synMuv suppressor genes encoding the chromatin remodeling and histone methylation factors *isw-1* and *mes-4* significantly suppresses the activation of *pals-5p::GFP* in the *lin-35(n745)* mutant under non-infection conditions. (A). Fluorescent micrographs of *pals-5p::GFP* (control) or *lin-35(n745); pals-5p::GFP* animals not exposed to Orsay virus or after Orsay virus infection. A brightfield image highlights the number of animals that were imaged. Scale bar is indicated. (B-D). Graphs showing a quantification of *pals-5p::GFP*(control) or *lin-35(n745);pals-5p::GFP* under RNAi of *isw-1* or *mes-4*. * p<0.001, NS: not statistically significant.

### *lin-15B* null mutants exhibit altered intestinal nuclear morphology

*C. elegans* infection with Orsay virus causes morphological changes in intestinal cells, including cell fusion, elongation of nuclei and nuclear degeneration (47). We looked for similar phenotypes using two strains that carry transgenes to visualize the intestinal nuclear membrane using the mCherry tagged nuclear pore protein NPP-9 (*ges-1p::npp-9::mCherry::BLRP::3xflag* or *ges-1p::npp-9::mCherry::BLRP::3xFLAG; his72p::birA::GFP*). One set of animals were infected with Orsay virus while the control group received no virus, and both groups were observed for 2 generations. The intensity of viral infection was established by the robust expression of *pals-5p::GFP* in animals carrying that transgene and infected simultaneously [**Fig 3. C, top panel]**. The no virus infection control animals had an average of 33.2 ± 2 intestinal nuclei at the adult stage. Orsay virus infected adults exhibited a 5% decrease in the number of intestinal nuclei, (31.5 ± 2.5) [**Fig 3. A & B**], weakly supportive of the previously observed nuclear degeneration, but not unassailable as a lone observation. We also observed elongation of intestinal nuclei in some Orsay virus infected animals [**Fig 3. C-middle panels].**

**Fig 3.**
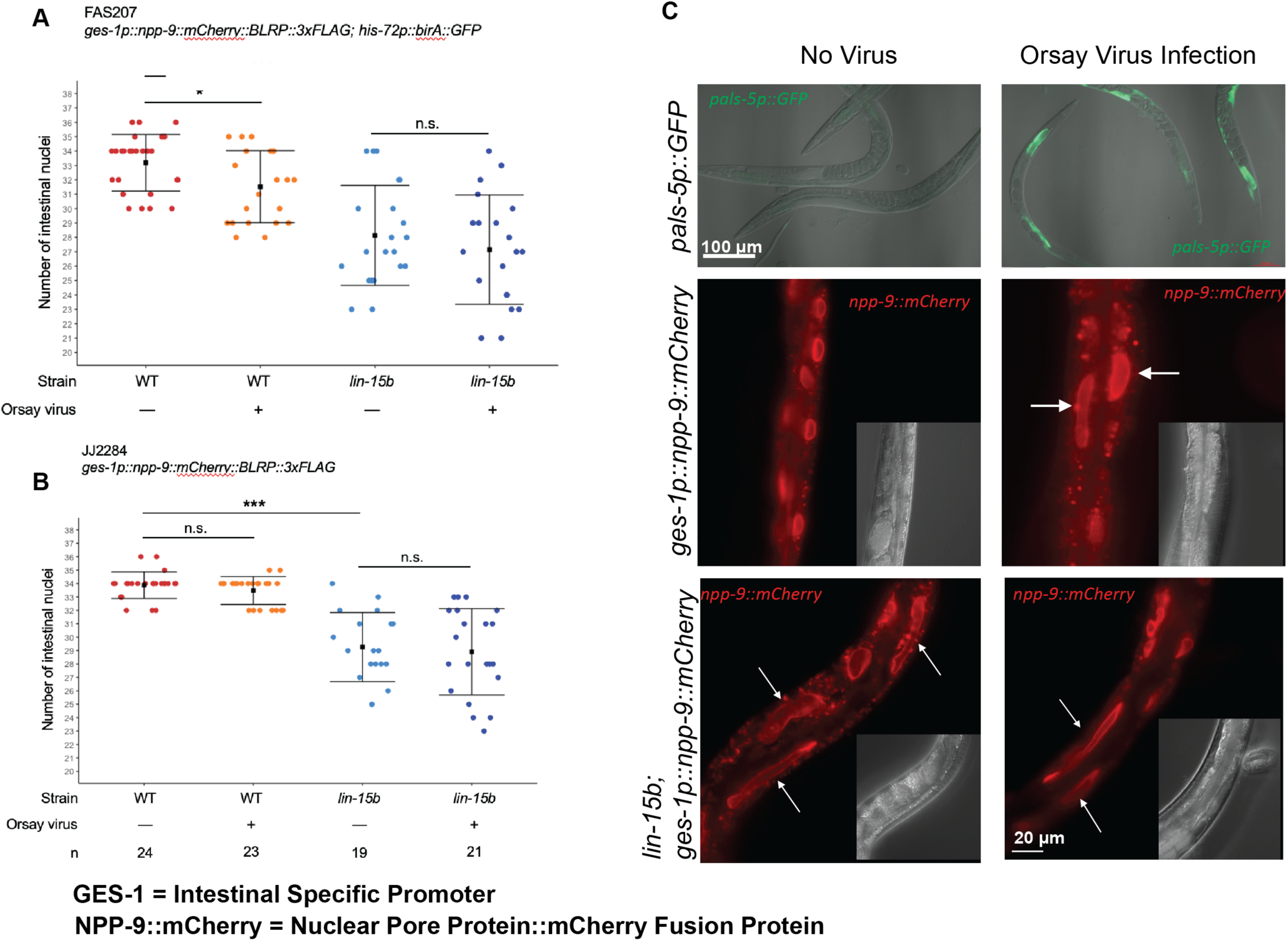
*lin-15b(-)* animals exhibit altered intestinal nuclear morphology. (A). Number of mCherry expressing intestinal nuclei in *FAS207* adults in wild-type and *lin-15b(W485*)* mutant backgrounds. (B). Number of mCherry expressing intestinal nuclei in *JJ2284* adults in WT and *lin-15b(W485*)* backgrounds. The square represents the average, and the bars represent the standard deviation. The sample size n has been indicated. p value calculated using ANOVA with Bonferroni correction. n.s: not statistically significant, * p<0.05, *** p<0.001. **C.** Representative images of *pals-5p::gfp* (top row), *FAS207* and *lin-15b(W485*)*; *FAS207* adults. White arrows indicate elongated intestinal nuclei. Scale bar is indicated. The graphs shown in A and B were generated using custom scripts in R studio and the statistical analysis were done using Jupyter Notebook.

To further test the intestinal nuclear response in synMuv B mutants to Orsay virus infection, we constructed a *lin-15B(W485*)* null mutant allele based on the canonical *lin-15B(n744)* null allele using CRISPR in the strain harboring the transgene *ges-1p::npp-9::mCherry::BLRP::3xflag*. We observed that under no virus infection conditions, *lin-15B(W485*)* animals exhibited a larger 15% decrease in the number of intestinal nuclei, with an average of 28.1 ± 3.5 nuclei [**Fig 3. A**]. L1 larval stage wild type animals normally have 20 intestinal nuclei which increase to 36 nuclei by the L2 stage, as most L1 intestinal cells undergo mitosis or at least nuclear division. Those 36 nuclei then increase their DNA content by undergoing DNA replication without cytokinesis during the next two larval stages and into adulthood, ending up with 36 nuclei with 32C genome copy number as roughly assayed by DAPI staining of DNA (34). Thus, the minimum number of nuclei we could expect to observe if there was no cell division at the first larval stage is 20; so, a decrease from 33 nuclei in wild type and to 28 nuclei in a *lin-15B* null mutant was substantial.

Strikingly, many of the intestinal nuclei in the *lin-15B(W485*)* animals were elongated [**Fig 3. C - lower panels**], suggesting this Orsay virus-associated cellular alteration is constitutively present in these *lin-15B (-)* animals. The decrease in the number of intestinal cells and the altered morphology of intestinal nuclei was also observed in a second independent line of *lin-15B(W485*)* that was generated in the *JJ2284* strain [See **S8 Fig**]. Orsay virus infection of the *lin-15B(W485*)* animals did not cause a further reduction of the number of intestinal nuclei or any change in the intestinal nuclear morphology. This implies that the intestinal nuclear response to viral infection of wild type *C. elegans* may be a direct result of downregulation of *lin-15B* or other synMuv B genes in Orsay virus-infected animals. Fusion of intestinal cells in Orsay virus infected animals has been reported previously (Felix et al, 2011). Both the observed decrease in the number of nuclei as well as the elongated appearance of the intestinal nuclei could be the result of a fusion of intestinal nuclei. Alternatively, we cannot rule out the possibility that the decrease in the number of intestinal nuclei in *lin-15B(-)* animals and Orsay virus infected animals could be due to a lineage defect or defect in karyokinesis during development.

### Orsay virus infection downregulates synMuv B gene activity to enhance antiviral RNAi

The *C elegans* Delta-like-ligand for the GLP-1 and LIN-12 Notch receptors is encoded by *lag-2. lag-2* is normally expressed only in the distal tip cells of the gonad and a subset of the vulval precursor cells (48). But mutations in several synMuv B genes cause nearly 100% penetrant ectopic *lag-2::GFP* expression in intestinal cells, which can be suppressed by gene inactivation of synMuv suppressor genes such as *isw-1* (14). If a *C. elegans* viral infection normally triggers the downregulation of synMuv B genes as a natural defense mechanism, we hypothesized that this may cause misexpression of *lag-2* within the intestinal cells as is observed in non-virally infected synMuv B mutants (14). We infected wild type animals carrying the *lag-2p::GFP* fusion gene with the Orsay virus and scored their F2 progeny for *lag-2p::GFP* misexpression in the intestine during the L4/adult stages. Upon infection of wild type animals with the Orsay virus, *lag-2p::GFP* was misexpressed in the intestinal cells with a 100% penetrance, similar to synMuv B mutants that are not infected with Orsay virus [**Fig 4A**]. This virus-induced intestinal misexpression of *lag-2p::GFP* was not suppressed by inactivation of synMuv suppressor genes such as *isw-1,* in contrast to the *lag-2p::GFP* induction in the synMuv B mutants [**S2 Fig. A&B**]. These data show that infection with Orsay virus recapitulates certain synMuv B loss of function phenotypes, for example, the induction of *lag-2p::GFP* in the intestine.

**Fig 4.**
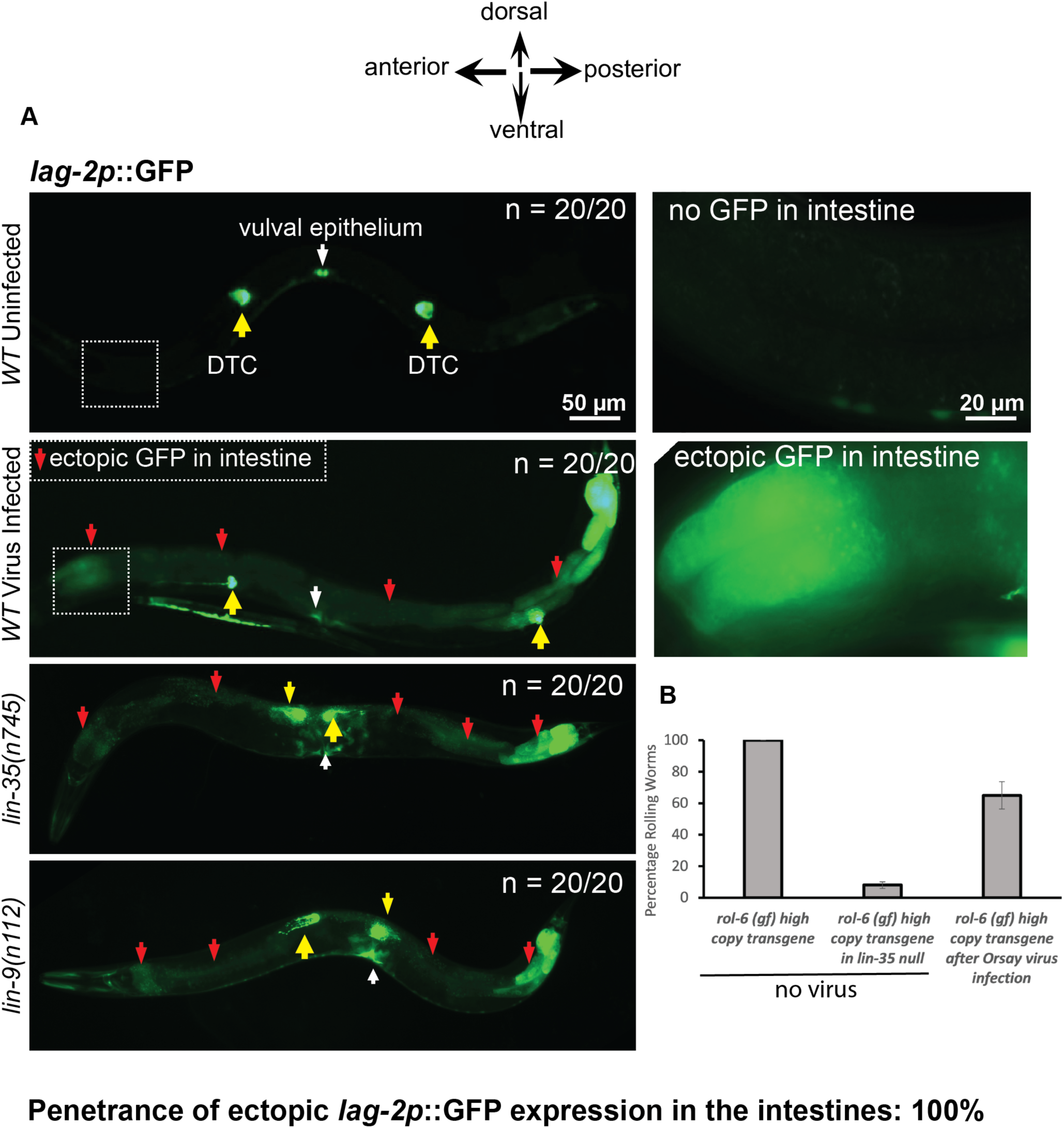
Orsay Virus infection causes misexpression of *lag-2p::GFP* in somatic tissues. (A). Shown are fluorescent micrographs depicting *lag-2p::GFP* expression in worms under no virus and under Orsay virus infection. The position of the distal tip cells of the germline are indicated by yellow arrows and the panels to the right of the WT/uninfected and WT/infected worms are panels of the region that is boxed on the left at a higher magnification. New ectopic expression of GFP seen within intestinal cells in the virus-infected animals is labelled with red arrows. Ectopic expression of GFP within intestinal cells of *lin-35(n745)* and *lin-9(n112)* mutants is shown as additional positive controls and the ectopic expression of GFP in the intestine is labelled with red arrows. The position of the central vulva is labeled with a white arrow. Scale bar is indicated. (B). Shown here is a quantification of the percentage of Rol phenotype of *mgIs30* animals that bear an integrated *rol-6* transgene, under no virus and infection with the Orsay virus. The suppression of the *rol-6* transgene in a *lin-35(n745)* null mutant is shown to provide a quantitative estimate of the intensity of the enhanced RNAi phenotype that is typical of most Eri mutants.

An integrated multicopy transgene (*mgIs30*) carrying a tandemly duplicated collagen mutant allele *rol-6(su1006),* an R71C substitution mutation of *rol-6* that causes the Rolling phenotype due to collagen assembly defects (49), causes a nearly 100% rolling (Rol) phenotype in wild type. But in nearly all Eri mutants (*eri-1, rrf-3, eri-6/7, lin-35, lin-15B*, and other synMuv B mutants), this transgene, like nearly all multicopy transgenes, is recognized as foreign and silenced so that the animals no longer roll (11). We reasoned that if downregulating synMuv B genes represent a natural part of the antiviral defense, then a viral infection itself may induce an Eri response, which may silence multicopy transgenes such as the *rol-6* transgene. We infected a wild-type strain carrying the *mgIs30* multicopy *rol-6* transgene with the Orsay virus and scored the percentage of rolling animals in the F2 generation. Consistent with our hypothesis, Orsay viral infection partially silenced the Rol phenotype of *mgIs30* [**Fig 4B]**, though to a weaker extent than what is observed in a synMuv B Eri mutant such as a *lin-35* null mutant, for example, where the silencing of the *rol-6* transgene is nearly 100% [**Fig 4B]**. This difference the level of transgene silencing may be due to the extent of inhibition of synMuv B gene activity, for example due to variations in viral titer or the synMuv B pathway response to a viral infection.

### Orsay virus infection of a *synMuv A* null mutant induce a Muv phenotype

Given that *lag-2p::GFP* is strongly misexpressed with a 100% penetrance within intestinal cells in wild type *C. elegans* infected with Orsay virus (which is strikingly similar to the *lag-2p::GFP* intestinal expression observed in synMuv B mutants), and the observation that Orsay virus infection in wild-type animals enhances RNA interference, we hypothesized that downregulation of some or all synMuv B genes may represent a normal step in viral defense. The simplest prediction of this hypothesis is that infecting a synMuv A mutant with the Orsay virus might down-regulate synMuv B activity as part of the normal response to a viral infection, so that, like a synMuv A; synMuv B double mutant, viral infection of a synMuv A mutant may cause a Muv phenotype. Only when <u>both</u> synMuv A and synMuv B genes are mutant does the multiple vulvae or Muv phenotype occur. If a viral infection in animals lowers synMuv B gene activity as a defense mechanism, we predicted that Orsay virus infection of a synMuv A null mutant might be sufficient to cause a Muv phenotype at least in some fraction of the population.

To test this, we infected the synMuv A *lin-8(n2731)* null mutant with the Orsay virus and examined F2 progeny for the appearance of the Muv phenotype. Strikingly, in 3 independent experiments, we found that about 10% of the F2 progeny were indeed Muv [**Fig 5A & B**], whereas uninfected *lin-8* mutant control animals did not produce any Muv progeny. The appearance of a Muv phenotype has never been reported in wild type *C. elegans* under Orsay virus infection conditions in any previous study nor in our own observations [**Fig 5B**].

**Fig 5.**
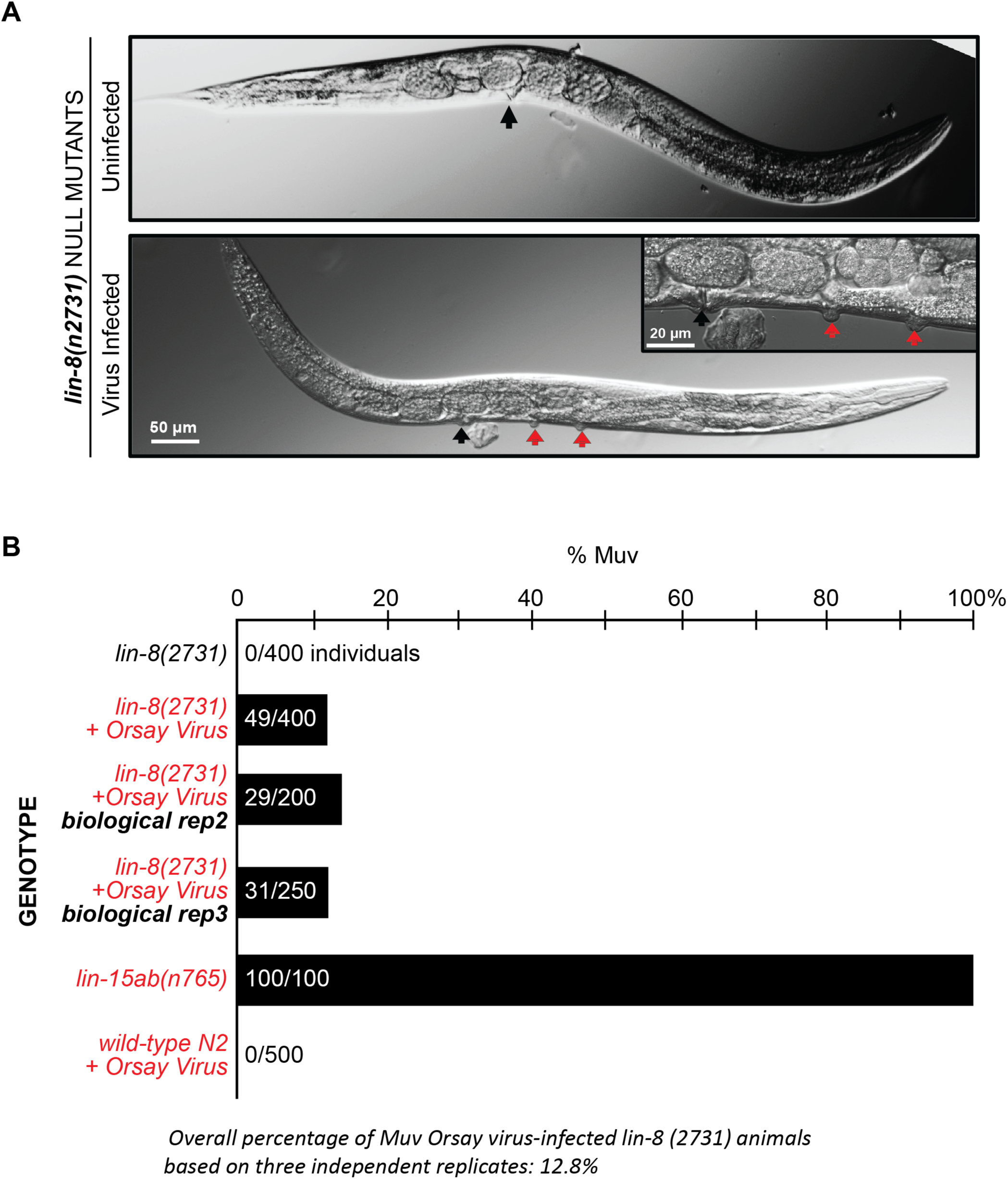
Orsay virus infection in *lin-8(n2731*), synMuv A mutants induces a Muv phenotype. (A). Brightfield micrographs showing uninfected vs virus-infected *lin-8(n2731)* mutant animals. The central vulva is indicated by a black arrow. The inset image in the panel showing the virus infected animal is a magnified region showing ectopic vulval inductions. The ectopic vulval inductions are indicated by red arrows. (B). Quantification of the percentage of Muv animals seen in *lin-8(n2731*) mutants under uninfected compared to Orsay virus infection. The Muv quantification was performed on the F2 generation of the virus-infected animals. Scale bar is indicated.

To follow the progeny of these Orsay infected synMuv A mutant animals, we singled out eight gravid *lin-8(n2731)* adult animals that were Muv after an Orsay virus infection, bleach treated them to remove any residual viral particles that may be present, and then allowed their eggs to hatch and examined their progeny at the adult stage to discern if the Muv phenotype persisted. We found that 100% of their progeny had a non-Muv vulval phenotype. We conclude that Orsay virus infection of *C. elegans* does not induce a mutation in a synMuv B gene in *lin-8(n2731)* null animals to give rise to a Muv phenotype.

Taken together, these findings suggest that downregulation of one or more synMuv B genes is a normal step in the antiviral defense response, and that viral infection induces an enhanced RNAi physiological state. The 10% penetrance of the Muv phenotype suggests that the downregulation of the synMuv B gene activity is just at the threshold for synergizing with the synMuv A mutations to confer a Muv phenotype (see Discussion).

### The siRNA landscape of synMuv B Eri mutants

Abundant endogenous siRNAs derived from thousands of genes are produced by wild type *C. elegans* that are not infected with viruses or exposed to specific dsRNAs that trigger additional exogenous RNAi of one targeted gene. These natural siRNAs regulate the mRNA levels and chromatin structures of a huge number of genes in wild type *C. elegans;* nearly all of these siRNAs are not produced in the RNAi-defective *mut-2* or *mut-16* mutants, and stunningly, these missing siRNAs from thousands of genes cause almost no phenotypes except for activation of normally dormant transposons and a failure to respond to injected or ingested dsRNAs (3, 4, 50, 51). In contrast, enhanced RNAi mutants disrupt the production of a far smaller subset of endogenous siRNAs. *C. elegans* siRNAs are classified by their length and 5’ nucleotide, which generally predict which of the nearly two dozen Argonaute proteins present them to specific mRNA and other target RNAs (52). The primary 26G siRNAs (26nt in length with G at the 5’ end), associate with the Argonaute proteins ERGO-1 and ALG-3/4. The targets of ERGO-1 26G siRNAs include about 100 protein coding genes, pseudogenes and long noncoding RNAs, many of which are integrated retrotransposons (51). The *ergo-1* mutants are strongly enhanced for RNAi, suggesting that either silencing these retrotransposons uses much of the RNAi bandwidth of wild type *C. elegans*, or that the viremia that results from failure to silence those retrotransposons with ERGO-1 siRNAs when *ergo-1* is defective, in turn enables a viral infection from those normally silenced retroviruses, to in turn induce an antiviral response, or increased RNAi, a sort of *C. elegans* inflammatory response. Targets of the ALG-3/4 26G siRNAs include spermatogenic genes (53–56); these 26G siRNAs and the Argonaute that mediate their recognition of mRNAs in sperm have not been implicated in normal RNAi or enhanced RNAi, And the 100x to 1000x more abundant 22G siRNAs (22nt in length with G at the 5’ end), are loaded into the many WAGO Argonautes, which are greatly expanded in the nematodes compared to other animals, and CSR-1. A large fraction of these 22G siRNAs is produced in the germline to mediate the massive level of gene silencing in the germline and to ensure that silenced genes from the parent continue to be silenced in progeny. CSR-1-associated 22G siRNAs are also expressed in the germline and modulate expression of germline genes as well as genes that mediate chromosome segregation at meiosis and mitosis (57, 58). The WAGO-associated 22G siRNAs target protein coding genes, pseudogenes and transposons (26). Almost all 22G siRNAs are depleted in the *mut-16* RNAi defective mutant, but most of these very abundant siRNAs are produced normally in the enhanced RNAi mutants *eri-6/7, ergo-1, rrf-3,* and *eri-1* (*10*). But the *eri-6/7* and *ergo-1* Eri mutants exhibit nearly complete depletion of the 26G siRNAs and a subset of secondary WAGO 22G siRNAs that are produced from these particular primary 26G siRNAs (12, 55). Because SynMuv B mutants exhibit a very strong Eri phenotype that is comparable to that of other well-studied Eri mutants, we investigated the landscape of small RNAs that are generated in these mutants.

Given that many of the key synMuv B mutant phenotypes are observed in polyploid somatic cells (the 8C hypoderm for LIN-3 activation and the 32C intestine for ectopic P-granule formation), we first carried out deep sequencing characterization of all siRNAs in synMuv B double mutants that produce little to no germline: *glp-4(bn2), lin-35(n745)*; *glp-4(bn2)* and *lin-15B(n744); glp-4(bn2)* grown at the non-permissive *glp-4* temperature of 25C [See **Fig 6 and S9 Fig & S10 Fig**]. The *glp-4(bn2)* mutation causes a temperature sensitive genetic ablation of the *C. elegans* germline when animals are grown at 25C. The complete list of the significantly upregulated and downregulated by a factor of 5-fold siRNAs are presented in the **Additional supporting information Table 1**.

**Fig 6.**
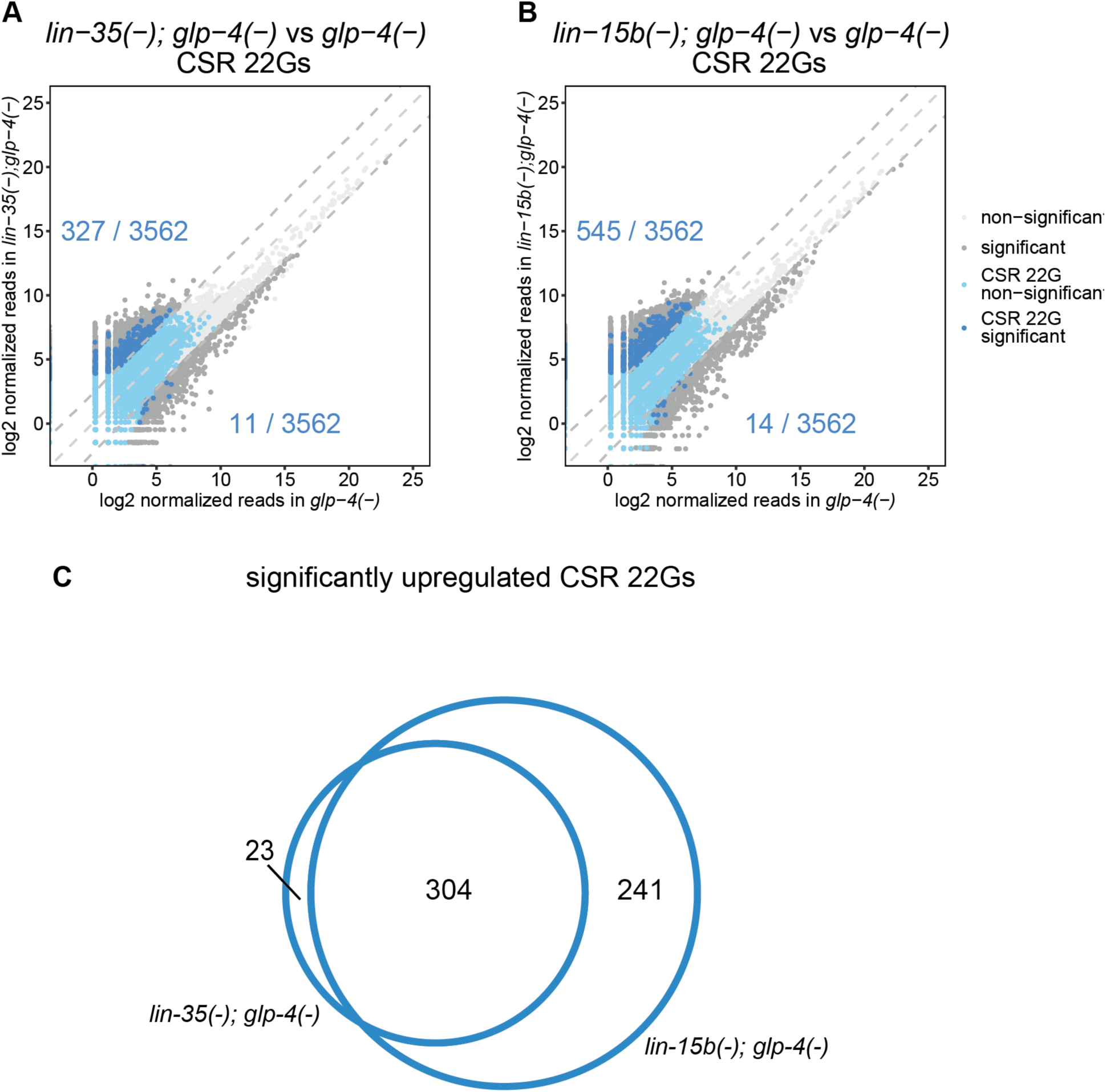
CSR 22Gs are upregulated in synMuv B mutants. **A-B.** Comparison of normalized reads of CSR 22Gs expressed in *lin-35(n745); glp-4(bn2)* (A) and *lin-15b(n744); glp-4(bn2)* (B) with *glp-4(bn2)*. Dark grey data points represent small RNAs that are differentially expressed by 5-fold and adjusted p value < 0.05. Dark blue data points represent CSR 22G siRNAs that are significantly upregulated by atleast 5-fold. Light blue data points represent other CSR 22G siRNAs. The rest of the small RNAs are represented as light grey data points. The lines denoting equal expression, 5-fold higher expression and 5-fold lower expression are shown as grey dashed lines. The dark blue numbers indicate the number of CSR 22Gs that are upregulated in the synmuv B backround (upper) or the control background (lower).**C.** Venn diagram representing overlap between CSR 22Gs upregulated in synmuv B mutants in (A) and (B).

We observed greater than a five-fold increase in about 10% of the detected CSR-associated 22G siRNAs in both *lin-35(n745)*; *glp-4(bn2)* and *lin-15B(n744); glp-4(bn2)* mutants compared to the *glp-4(bn2)* control animals [**Fig 6 A & B**]. There was significant overlap between the genes targeted by the upregulated CSR-22Gs in both *lin-35(n745)*; *glp-4(bn2)* and *lin-15B(n744); glp-4(bn2)* mutants, suggesting that a common pathway is misregulated in these distinct synMuv B mutants [**Fig 6C**]. SynMuv B mutants are also known to misexpress many genes that are normally germline-restricted in somatic tissues such as the intestine (Wang et al, 2005, Petrella et al, 2011). Because the animals that we analyzed have no germline, the significant increase in CSR-22G siRNAs that are observed in these synmuvB mutants may reflect the misexpression of these germline genes in the intestine. We also observed loss of about half of the detected ERGO-26G siRNA in both *lin-35(n745)*; *glp-4(bn2)* and *lin-15B(n744); glp-4(bn2)* **[See S9 Fig. C and S10 Fig. C]**. This partial loss of ERGO-26G siRNAs in synmuvB mutants is less severe than the complete depletion of ERGO 26G siRNAs in *eri-1*, *eri-6, eri-7* and *ergo-1* loss of function mutants (12, 55). It is possible that the synMuv B mutants affect 26G siRNA production in the intestine for example, whereas other *eri-*mutants may affect more cell types.

WAGO-22G siRNAs were differentially expressed in both *lin-35(n745)*; *glp-4(bn2)* and *lin-15B(n744); glp-4(bn2),* with some showing increased and others showing decreased expression [**See S9 Fig. G, S10 Fig. G]**. We also observed a decrease in the expression of a few microRNAs and piRNAs in both *lin-35(n745)*; *glp-4(bn2)* and *lin-15B(n744); glp-4(bn2)* [**See S9 Fig. K, L and S10 Fig. K, L].** An exciting possibility is that these particular miRNAs, for example *mir-8191* may be located, in locally amplified regions that depend on the dREAM complex, as has been observed for the Drosophila chorion genes (35–38). Finally, we found very little change in the abundance of ALG class 26G siRNAs [**See S9 Fig. A, S10 Fig. A]**.

Additionally, we carried out in parallel, the deep sequencing of small RNAs in *lin-15B(n744), lin-35(n745), lin-9(n112)* and *lin-52(n771)* synMuv B mutants with intact germlines. Here, we did not find any striking differential expression of any of the classes of small RNAs, possibly because the abundance of small RNAs derived from the germline may mask the changes in siRNA expression that may occur in the soma [**See S11-S14 Fig**].

### Molecular epistasis analysis of the enhanced RNAi in synMuv B mutants

The 26G siRNAs in animals are synthesized via the action of the RdRPs RRF-1,2,3 and EGO-1, which produce longer dsRNAs that are further processed into smaller 26G-RNAs through DICER and PIR-1/phosphatase (54–56, 59, 60). Our deep sequencing analysis of the synMuv B mutants is consistent with the decline in 26G siRNAs in most Eri mutants,such as *eri-1(mg366*) (55) or *eri-6/7* and *ergo-1* which with dramatic declines in 26G siRNAs (10, 12, 61). To further explore the 26G axis of siRNAs, we mined mRNAseq datasets (from the NCBI Geo-collection) of *lin-35(n745)* and *lin-15B(n744)* mutants, to see whether transcripts of any genes that are known to cause enhanced RNAi (such as *eri-*genes, *rrf-3* etc), when mutant were downregulated in synMuv B mutants. However, we did not observe lower mRNA levels for any of the known genes that mutate to an enhanced RNAi phenotype in the *lin-15B* and *lin-35* mRNA-seq datasets [**See Additional supporting information data tables 2-4**]. This suggests that the decrease in 26G siRNAs in the synMuvB mutants is not due to the downregulation of other known *eri-*genes. 26G-siRNAs are bound by ERGO-1 Argonaute which then targets those siRNAs to particular target mRNAs, approximately 100 recently acquired retroviral elements (10–12). To test whether synMuv B mutations affect ERGO-1 protein levels or localization to in turn affect 26G siRNA levels, we observed the ERGO-1 protein using a GFP-tagged full length ERGO-1 protein (*tor147[GFP::3XFLAG::ergo-1*, the longest predicted isoform (62). We used RNAi to inactivate synMuv B genes or to inactivate genes that encode suppressors of synMuv B genes, and then observed the localization and expression of *GFP::3XFLAG::*ERGO-1 to determine if downregulating the activity of either synMuv B genes or synMuv Eri suppressor genes can impact the spatiotemporal expression of GFP tagged-*ergo-1* in animals. The GFP tagged-ERGO-1 fusion protein was strongly expressed in the seam cells during the L3 stage and later in the vulval epithelial cells in the mid-L4 and young adult stages [**S3 Fig. A-F**]. We targeted the synMuv B genes *lin-9* and *lin-13* [**S3 Fig. G-N**], and the synMuv suppressor gene *isw-1* by RNAi [**S3 Fig. O-R**] and found that the *ergo-1* expression pattern was unchanged. Thus ERGO-1 is unlikely to be a direct or indirect target of synMuv B protein regulation.

### The expression pattern and subcellular localization of *lin-15B* and *lin-35* synMuv B proteins

A detailed description of the subcellular expression/localization patterns for any synMuv B gene product has not been previously reported. We investigated the expression patterns and subcellular localization of the LIN-15B and LIN-35 proteins using *LIN-15B::EGFP* and *LIN-35::EGFP* protein fusions, both generated by Mihail Sarov via a recombineering pipeline (63). In these fusion protein reporter strains, the synMuv B gene is tagged by a protein fusion to EGFP at its C terminus in the context of roughly 40kb of genomic DNA cloned on a fosmid, and then subsequently biolistically transformed at a relatively low copy number (63). These strains are likely to recapitulate the endogenous expression/localization patterns of their tagged proteins and given their low copy number representation, they may not be subject to the transgene silencing typical of high-copy number transgenes that is commonly associated with Eri-genotypes. These synMuv B GFP fusion proteins were previously used to map the binding sites of the synMuv B proteins on the promoters of genes genome wide (64). We detected robust LIN-15B::EGFP protein localization in both the hypodermal cells and intestinal cells during the mid-L4 stage and later animals. Intriguingly, LIN-15B::EGFP is localized subnuclearly, forming punctate EGFP foci in the hypodermal and intestinal nuclei. These foci may represent genomic binding sites for client genes of the LIN-15B THAP domain transcription factor. To quantitate the abundance of LIN-15B::EGFP, we quantitated GFP in particular hypodermal cells and intestinal cells from the anterior, middle, and posterior [**S4 Fig**]. We measured LIN-15B::EGFP fluorescence intensity in these cells using Image J and the number of LIN-15B::EGFP subnuclear foci was manually scored.

These subnuclear foci of LIN-15B do not change under RNAi targeting genes such *mut-16*, *rde-1* or *rde-4*, which when deactivated impair RNA interference [**See S5 Fig**]. The subnuclear foci of LIN-15B also remain unperturbed under RNAi that targets genes known to enhance RNAi such as *eri-6, lin-9, lin-13, lin-35, lin-37* and *lin-52 or* genes that are known to suppress the enhanced response to RNAi of synMuv B mutants such as *isw-1* and *mes-4* [**See S5 Fig**]. Crossing lin*-35(n745)* null mutant into the LIN-15B::EGFP reporter strain did not change LIN-15B localization or the average number of subnuclear foci [**See S6 Fig. A-D**]. Taken together, these findings suggest that the LIN-15B localization is not coupled to its role to activate the antiviral RNAi response.

Similar observations of the *lin-35*/retinoblastoma expression pattern and protein localization using a *lin-35::EGFP* translational fusion gene revealed that LIN-35::EGFP is strongly expressed in all intestinal cells, where it localizes to the nucleus as well as to the nucleolus [**S7 Fig. A & B**]. Each intestinal cell showed one or two LIN-35::EGFP nucleolar inclusions as well as uniform localization across the nucleus [**S7 Fig. A & B**]. There were no obvious changes to LIN-35 localization or expression level in strains induced to have an enhanced or defective RNAi by inactivation of *eri* or *mutator/rde* genes [**S7 Fig. C, D & E**].

## DISCUSSION

The potency of an injected, in vitro transcribed, 1 kb dsRNA for *C. elegans* RNA interference, discovered by Fire and Mello almost 30 years ago, was the tip of a very large RNAi iceberg. In fact, the complexity of the endogenous small RNA world immediately emerged from the genetic analysis of RNAi: the first RNAi-defective mutants identified included a large set of known mutator genes that had already emerged from searches for mutations that enhance transposition or reversion of transposons; defects in RNA interference was the molecular basis of many mutator mutants that release transposons from RNAi control (1, 4, 5, 7, 65, 66). Mutants that disable RNAi activate transposable elements, integrated retroviruses in the *C. elegans* genome, the remnants of past successful retroviral infections that are normally silenced by RNAi, which now become active if key RNAi components are defective to then insert or revert from an insertion in any gene (65). Thus, RNAi is not just a response to artificial injected dsRNAs; there are natural endogenous dsRNAs that produce siRNAs which the same genetic pathways mediate their activities.

Transposons are retroviral elements that have been thought to have successfully evaded surveillance and integrated into the genome. They are universal to eukaryotes, with an even higher fraction in the mammalian genome (67) than *C. elegans*, perhaps protected by its more elaborate RNAi pathway, most especially the RNA-dependent RNA polymerase siRNA amplification sub-pathway. These transposable elements integrated in genomes are the seeming victors from the battle between host RNA interference and invading viruses. But these integrated retroviruses are perhaps not viral trophies to their past victories over the host genome, but rather they may constitute host ammunition for future defense of newly evolved but related viruses that are likely bear sequence similarity to the viral carcasses left behind in the genome. Those retroviruses that have evaded defense to become integrated into host genomes may not have actually escaped RNAi immunity; the highly evolved and ramified RNAi pathways of *C. elegans* continue to actively but reversibly repress their activity, perhaps to allow their re-activation during a triggering of defense of new viruses. For example, the endogenously produced siRNAs that silence integrated retroviruses may show some mRNA silencing activity on infection by new viruses which are related to the integrated viruses or the release of repression of integrated viruses to allow a “programmed” viral infection, may activate antiviral cascades that confer immunity to any virus. The early discovery from the genetic analysis of RNAi that the RNAi pathway has a major role in the reversible silencing of these viruses hints that there may be physiological states where their silencing is reversed, states where transposon hops are selectively advantageous. In addition, reversibly silenced viruses are a useful mechanism to program mutagenesis by their release from repression when it is needed. It is not a coincidence that the first RNAi defective mutations in *C. elegans* had already been identified as mutator gene, with unleashed transposons for generation of genetic diversity (3, 68). For example, under conditions of viral or bacterial pathogen attack, the production of new mutations by the transposition of transposons may be selectively advantageous. Thus, these retrotransposons may have been domesticated by host RNAi to only allow their transposition and thus production of genetic variation by mutagenesis when variation is advantageous.

Similarly, the *C. elegans* enhanced RNAi mutants have highlighted a similar physiological regulation of antiviral defense or transposon mobilizations; enhanced RNAi or Eri mutants have emerged from genetic screens at about the same frequency as RNAi defective mutations (Rde, Mut, etc). Each of the mutant classes, RNAi defective (Rde) and enhanced RNAi (Eri), points to these phenotypes as physiological states, with the mutants frozen in a state of either no RNAi defense or higher than normal RNAi defense. Conversely, *Saccharomyces cerevisiae* jettisoned its RNAi pathway to allow its permanent infection by the killer RNA virus (69).

The synMuv B genes emerged from large genetic and RNAi screens for enhanced RNAi mutants or gene inactivations (7, 11), but were discovered 20 years earlier in unrelated comprehensive genetic analysis of receptor tyrosine kinase signaling during *C. elegans* vulval development (70, 71). Our analysis of the mRNA expression profiles of synMuv B null mutants showed a significant overlap between genes upregulated in the synMuv B mutants and those induced upon viral infection by the Orsay RNA virus (40). Notably, the 100x or more induction of genes in synMuv B mutants that are also strongly induced by Orsay virus, for example, members of the PALS gene family, suggests a potential link between synMuv B mutations and the activation of a coordinated genetic response akin to that observed during viral infections. The dramatic upregulation of Orsay virus response genes in synMuv B mutants, even in the absence of viral infection, suggested that the down-regulation of synMuv B gene activity could mimic a physiological state of viral resistance, enhanced RNAi, that is normally induced in an actual viral infection, to activate antiviral defense mechanisms, including RNAi. The robust induction of *pals-5p::GFP* expression in synMuv B mutants, even in the absence of actual viral infection, suggests that these mutants constitutively activate their antiviral defense pathways.

Because the *C. elegans* synMuv B mutants *lin-35* and *lin-15B* activate highly congruent gene expression responses to those of an Orsay RNA virus infection, we tested whether an actual Orsay infection down-regulates synMuv B activity: whether an Orsay virus infection can induce a Muv phenotype in a synMuv A mutant, just as a synMuv B; synMuv A double mutant shows the Muv phenotype. Satisfyingly, we found that Orsay viral infection of a synMuv A mutant causes a reproducible 10% penetrant Muv phenotype, but Orsay viral infection of wild type animals does not cause a Muv phenotype. This supports the hypothesis that the downregulation of one or more synMuv B gene activities is a key defense response during a viral infection, and that a viral infection induces a physiological state that is like the enhanced RNAi state of a synMuv B mutant even in the absence of a viral infection. Why is the penetrance of the Muv phenotype not higher than 10% during a viral infection of a synMuv A mutant, when null mutations in synMuv B in the background of a null synMuv A mutation can cause 100% penetrant Muv phenotype? The most likely explanation is that a viral infection lowers synMuv B activity but not to zero; that it is most similar to a non-null synMuv B mutant with a null synMuv A mutant. In addition, the Muv phenotype of a synMuv A; synMuv B double mutant is caused by the loss of the synMuv B gene specifically in the hypodermis; the loss of synMuv B in the hypodermis is specifically required for the Muv phenotype of a synMuv A; synMuv B double mutant (22). The loss of synMuv B and synMuv A in the hypodermis causes more than 100x increased expression of the LIN-3 ligand which is secreted to the adjacent affect vulval cells to cause the Muv phenotype in those vulval cells (15). Because Orsay virus infection induces *pals-5p::GFP* in the intestine, the primary tissue where the virus infection is thought to occur via its ingestion (41), it is possible that synMuv B down-regulation in the hypodermis during a viral infection of the intestine is far less than a synMuv B null mutant, thus the lower penetrance of the Muv phenotype.

We also explored the link between gene inactivations that suppress the enhanced RNAi in synMuv B mutants and their activation of antiviral response genes. RNAi inhibition of known suppressors of the enhanced RNAi phenotype of synMuv B genes, such as *isw-1* and *mes-4*, caused a significant reduction in *pals-5p::GFP* induction by a synMuv B mutation, indicating a potential interplay between RNAi and antiviral defense pathways regulated by synMuv B genes. Interestingly, our findings suggest that the Eri phenotype induced by synMuv B mutations may engage distinct pathways from other known Eri mutants, such as mutations in the RNA dependent RNA polymerase *rrf-3* or the *eri-1* exonuclease, highlighting the complexity of antiviral defense mechanisms in *C. elegans*. The observed suppression of *pals-5p::GFP* expression upon *isw-1* and *mes-4* knockdown in synMuv B mutants under non-infection conditions underscores the role of these genes in modulating the antiviral response. However, inactivation of *isw-1* or *mes-4* did not suppress the induction of *pals-5p::GFP* during an actual viral infection, suggesting that synMuv B induction of the *pals-5p::GFP* response, may be one of several pathways that regulates the expression of *pals-5* during an actual viral infection.

*C. elegans* RNAi pathways surveil thousands of genomic loci in wild type *C. elegans* under non-viral infection conditions to produce hundreds to thousands of siRNAs per million reads. Most of this massive cloud of siRNAs against their own genes are missing in animals with defects in RNAi, most dramatically in the Mutator mutants (Wallis et al., 2019). In contrast, null mutations in the enhanced RNAi mutant *eri-6/7*, encoding an RNA helicase, disrupts the production of a far more circumscribed set of siRNAs that silence approximately 100 integrated retroviral elements (10–13). This Eri mutant may be enhanced for RNAi because the desilenced retroviral elements trigger ER stress, probably due to the replication of viral sequences in the ER, and the enhanced RNAi of an actual or falsely sensed infection (10).

Our survey of the small RNA landscape in synMuv B Eri mutants showed a 5-fold increase in 10% CSR-class 22G siRNAs, 5-fold decrease in 50% of ERGO-class 26G siRNAs, 7% of WAGO-class 22G siRNAs, 4% of piRNAs and 14% microRNAs in synmuvB mutants. The partial loss of ERGO-26Gs and piRNAs in synmuvB mutants is distinct from the previously characterized *eri* mutants (12, 55). One model for the molecular basis of enhanced RNAi mutants is that there is a limiting component to RNA interference; for example, the silencing of transposons uses some limiting factor for RNAi such that if the silencing of retrotransposons is disabled, that limiting factor is released for the dsRNA-programmed silencing of feeding RNAi (61). However, the differences in the small RNA landscape of the synMuv B mutants suggest a distinct mechanism may underlie the enhanced RNAi of synMuv B mutants: the targets of ERGO-class 26Gs, piRNAs and WAGO-class 22Gs include pseudogenes, foreign elements and transposons, loss of these small RNAs in synmuvB mutants may cause the misexpression of these genes and may explain the constitutively active anti-viral readiness of the synmuvB mutants. It is possible that the de-repression of integrated viruses, the activation of transposons, in the synMuv B mutants in fact causes a viremia of those viruses to in turn induced stronger RNAi. The induction of enhanced RNAi during a viral infection makes excellent biological sense.

The synMuv B genes encode homologues of the dREAM complex, which are conserved across animals and plants, including the LIN-35/retinoblastoma (Rb) tumor suppressor, and the Rb complex components LIN-53 /RbAp48, LIN-37/MIP40, LIN-54/MIP120, DPL-1 /DP, LIN-9/MIP130 (16, 19–21). The synMuv A genes *lin-8, lin-56*, and *lin-15A* encode widely expressed nuclear proteins with no obvious homologues outside of nematodes (17, 18), and in the case of *lin-8*, there exists more than a dozen paralogues in *C. elegans* and many dozen in *C. japonicum*, located in clusters of duplicated paralogues (45). The nuclear localization of the synMuv A proteins is consistent with their strong genetic interaction with the nuclearly localized synMuv B dREAM complex proteins, but nothing in the anonymous synMuv A protein sequences, even though they encode nuclear proteins, suggests their mechanism of action.

While the analysis of the synMuv A and synMuv B mutants in specification of the ventral precursor cells has been extensive and beautiful, how it connects to the enhanced RNAi of the synMuv B mutants is not instantly obvious from the beautiful molecular genetic analysis of LIN-3 signaling (15, 16, 20, 21, 23). The molecular basis of how the DNA-binding proteins and associated proteins of the synMuv B pathway actually intersect with the PIWI proteins and RdRPs that are central to RNAi has not been characterized. Our investigations suggested that RNAi suppression of the Eri-phenotype of synMuv B mutants by *isw-1* or *mes-4* RNAi do not affect the localization of LIN-15B or LIN-35 (See Supplemental Figures 5 & 7). This suggests that a molecular mechanism distinct from the localization or abundance of the SynMuv B proteins regulates the intensity of the RNAi pathway.

However, extensive analysis of the many synMuv B orthologues in Drosophila has revealed their activity in endoreduplication of particular client genes and the detailed analysis of *C. elegans* synMuv B genes in the vulval cell signaling pathway has revealed an analogous intersection with endoreduplication. The Drosophila dREAM complex binds specifically at replication origins that flank the Drosophila chorion genes to control their local (about 50kb) endoreduplication to nearly 100x the gene dosage of the normally diploid Drosophila genome (35). This enables the regulated production of the abundant eggshell proteins; mutations in the dREAM complex cause a humpty dumpty phenotype, fragile eggs due to insufficient production of chorion proteins (37, 72). The dREAM complex proteins bind directly to these chorion gene replication origins and are necessary for the localized amplification of just those genes; in the Drosophila retinoblastoma mutant, unregulated endoreduplication occurs (36–38).

This intersection of the dREAM complex with endoreduplication is also a hallmark of the synMuv B gene function in LIN-3 growth factor signaling from the polyploid *C. elegans* hypodermis to the adjacent vulval cells that show the Muv phenotype in the synMuv B mutants. The polyploid hypodermal cells hyp7 cell is the focus of synMuv B gene function in *C. elegans* for the regulation of LIN-3 production (15). The *C. elegans* hypoderm is normally endoreduplicated to 8C (3 full genome duplications) or more at adult stages (73).

Analysis of other shared *C. elegans* synMuv B mutant phenotypes also suggests that they also function in another polyploid tissue, the intestine, to control the production of P-granules which are approximately 1 micron ribonucleoprotein granules intimately connected to RNAi (29). The P-granules are normally germline-specific multi-protein and RNAs that assemble into the originally described LLPS particles of the single blastomere that is the precursor of the *C. elegans* germline (27). Loss of function mutations in multiple *C. elegans* synMuv B genes cause misexpression of these normally germline-specific P-granules in somatic tissues, most dramatically in the polyploid intestine and hypodermis (7, 28). The adult *C. elegans* intestine is normally endoreduplicated from diploid at hatching to 32C (5 inferred full genome duplications; local endoreduplications not yet studied with the DAPI-light microscope DNA quantitation), with one duplication at each larval stage but no chromosome condensation or mitosis (25). The hyp7 hypoderm where ectopic P granules are also observed is 8C. The P granules, as macromolecular complexes that are visible at light microscope magnification, are filled with abundant proteins and RNAs, which like chorion proteins, represent candidate genes for localized endoreduplication (28, 30–33, 74).

It may be significant that the *pals* genes as well as the *lin-8*-like genes (*lido-genes*) localize to genomic clusters with several *pals* or *lido* genes within cluster. The clusters are also very plastic in *Caenorhabditis* evolution, varying down to just 3 *lido* genes in *C. remaniae* and up to 58 *lido* genes in *C. japonicum*. These clusters of poorly conserved *pals* and *lido* genes may be under dREAM complex control of their local amplification, which may serve to increase the gene copy number for example in the intestine.

That RNAi is highly regulated is not surprising: RNAi has evolved by selection for potent antiviral defense during almost a billion years of eukaryotic infection by viruses. But it is surprising that the dREAM complex is so central to RNAi. And the mammalian-conserved synMuv B pathway reveals new pathways to activate production or response to small interfering RNAs and may be targeted with drugs to activate human antiviral responses or responses to RNAi-based pharmaceuticals.

## MATERAILS AND METHODS

### Nematode Strains Used

#### Strains made available through the CAENORHABDITIS GENETICS CENTER (CGC)

*N2 (wild-type), lin-35(n745), lin-52(n771), lin-9(n112), lin-15B(n744), lin-15AB(n765), eri-6/7(mg441), eri-1(mg366), NL2099 [rrf-3(pk1426], ERT54 [jyIs8 [pals-5p::GFP; myo-2p::mCherry], mgIs30 [rol-6(su1006)::lin-14 3′-UTR, col-10::lacZ, lim-6::gfp]), OP184[wgIs184 [lin-15B::TY1::EGFP::3xFLAG + unc-119(+)], OP763[wgIs763 [lin-35::TY1::EGFP::3xFLAG + unc-119(+)], lin-8(n2731), qIs56(lag-2::gfp, unc-119(þ), eri-1(mg366), eri-6(mg379), rrf-3(pk1426)*.

More details regarding the gene mutations featured here can be found at http://www.wormbase.org

#### Strains generated in our lab for this study

*lin-35(n745) glp-4(bn2), glp-4(bn2); lin-15B(n744), glp-4(bn2), lin-35(n745); jyIs8, lin-52(n771); jyIs8, lin-15B(W485*), lin-15B(W485*); jj2284, lin-15B(W485*);FAS207, eri-1(mg366); jyIs8, eri-6(mg379); jyIs8, rrf-3(pk1426); jyIs8, lin-9(n112); qIs56, lin-35(n745); qIs56*.

#### Strains sent by others*

*FAS207*, *unc-119(ed3) III; zuIs261[ges-1(5’UTR)::npp-9::mCherry::BLRP::3xFLAG::npp-9(3’UTR); unc-119(+)]; zuIs236[his-72(5’ UTR)::BIRA::GFP::his-72(3’ UTR); unc-119(+)]*.

JJ2284 *unc-119(ed3) III; zuIs261[ges-1(5’UTR)::npp-9::mCherry::BLRP::3xFLAG::npp-9(3’UTR); unc-119(+). *These strains were kindly provided by Steven Henikoff and Florian Steiner*.

#### Generating the strain *lin-15B(W485*)*

In brief, gravid animals were injected with a CRISPR/Cas9 mix containing 30pmol *S. pyrogenes* Cas9 (IDT), 90pmol tracrRNA (IDT), 95pmol crRNA oHG6, 2.2mg repair template oHG8, and 80ng of *PRF4::rol-6 (su1006)* plasmid. The F1 progeny that exhibited the *rol* phenotype as well as their siblings were singled and genotyped for the edit using primers oHG9 and oHG10, followed by a restriction digest using AseI (New England Biolabs).

### Oligos Used

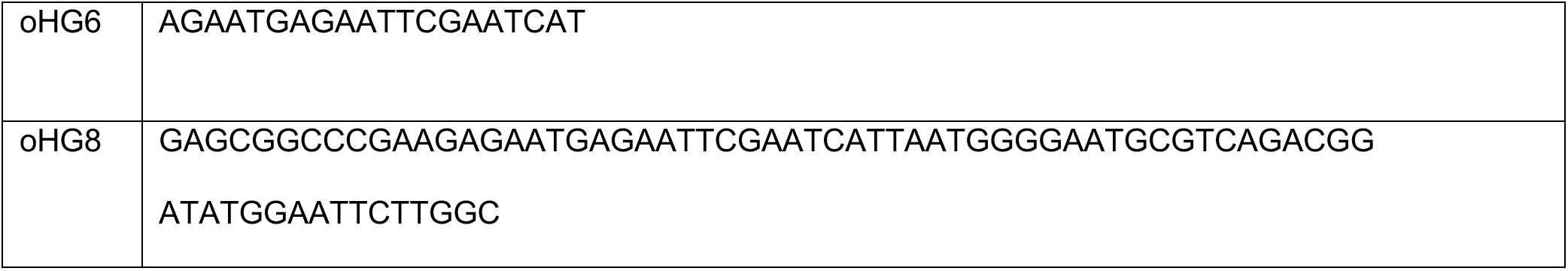
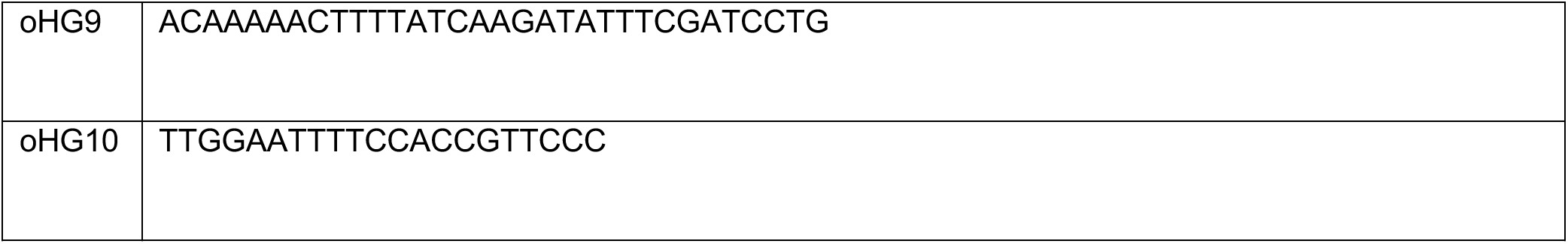

#### Counting intestinal nuclei

Animals infected with Orsay virus were collected in M9, crushed and filtered through a 22mm filter. 50uL of the resulting lysate was used to infect animals. Equal volume of M9 was used for the mock infection. Four L4 animals were infected with mock or Orsay virus lysate. Animals were allowed to propagate for 2 generations. When the animals were running out of food, the animals were chunked to fresh plates pre-infected with the Orsay virus lysate. F2 adults were picked into sodium azide on an agarose pad and imaged on the Zeiss Axio Imager Z.1 within an hour. The number of mCherry expressing intestinal nuclei was scored using the 40X objective.

#### siRNA Sequencing: Growth Conditions

Eggs were hatched and arrested as synchronous L1 larvae and then plated on *E. coli* OP50 and grown at 25°C until most had reached gravid adult stage (72-74 hours). At this time point, there was some asynchrony within the individual strains because of differences in development.

#### RNA Isolation

RNA was isolated from whole animals using Trizol and chloroform extraction according to the manufacturer’s recommendations but with the addition of a second chloroform extraction step (Life Technologies, cat# 15596018).

#### Small RNA Sequencing

Total RNA was treated with RppH to reduce small RNA 5’ triphosphates to monophosphates following the manufacturer’s recommendations (New England Biolabs, cat# M0356S)(75). 16-30-nt small RNAs were size selected on a 17% polyacrylamide/urea gel and purified using the ZR small-RNA PAGE Recovery Kit (Zymo Research, cat# R1070). Small RNA sequencing libraries were prepared using the NEBNext Multiplex Small RNA Library Prep Set for Illumina following the manufacturer’s recommendations but with the 3’ ligation step changed to 16 °C for 18 hours to improve capture of methylated small RNAs (New England Biolabs, cat# E7300S). PCR products corresponding in size to adapter-ligated small RNAs were size selected on a 10% polyacrylamide non-denaturing gel, electrophoretically transferred to DE81 chromatography paper, eluted at 70°C for 20 minutes in the presence of 1 M NaCl, and precipitated at -80°C overnight in the presence of 13 ug/ml glycogen and 67% EtOH. Small RNA read processing, including adapter trimming, quality filtering, mapping, and counting, was done with tinyRNA v1.5 using the default configuration (52). Reference sequences and annotations were based on the *C. elegans* WS279 release (76).

### Generating the top 100 Up and Down list of siRNA targets

- DESeq2 analysis using R was performed on three sample groups at once (i.e., glp*-4; lin-15B* and *lin-35 glp-4* and *glp-4*) for the 0mm (mismatch) set.
- Pairwise comparisons (i.e., *lin-15B; glp-4* and *glp-4* OR *lin-35; glp-4* and *glp-4*) yielded log2FoldChange and adjusted p-values (padj). To identify the siRNAs with significant changes ***up*** in the synMuv B mutants, the list of siRNA targets was filtered as follows:

`o` Greater than or equal to 5-fold change (i.e., log2FoldChange = log2(5) = 2.32);
`o` Less than or equal to adjusted p-value of 0.05; and
`o` Greater than or equal to 10 counts of normalized reads in the sy*nMuv B; glp-4* strains.
- To identify the siRNAs with significant changes ***down*** in the synMuv B mutants, the list of siRNA targets was filtered as follows:

`o` Less than or equal to 0.2-fold change (i.e., log2FoldChange = log2(0.2) = -2.32);
`o` Less than or equal to adjusted p-value of 0.05; and
`o` Greater than or equal to 10 counts of normalized reads in the *glp-4* strains.
- Each of the lists of siRNA targets were arranged in the order of greatest fold change (i.e., greatest log2FoldChange for siRNA targets that are the most upregulated and lowest log2FoldChange for siRNA targets that are the most downregulated).

### mRNA sequencing analysis

Fastq files were downloaded from GEO (GEO Accession numbers GSE62833 and GSE155190). The STAR aligner (77), was used to map sequencing reads to transcriptome in the ce10 reference genome. Read counts for individual genes were produced using the count function in HTSeq (78), followed by the estimation of expression values and detection of differentially expressed transcripts using EdgeR (79), which included only the genes with count per million reads (CPM) > 1 for two or more samples. Differentially expressed genes were defined by the cutoffs of > log2 fold change and false discovery rate (FDR) < 0.05. Volcano plot was generated using plotly and dplyr (80) in R.

### GEO accession numbers and genotype information

*GSM4697084 (N2 starved L1-rep1), GSM4697085 (N2 starved L1-rep2), GSM4697089 [lin-35(n745) starved L1-rep1)], GSM4697090 (lin-35(n745) starved L1-rep2), GSM4697102 (lin-15B(n744) starved L1-rep1), GSM4697105 (lin-15B(n744) starved L1-rep2), GSM1534086 (lin-35[JA1507(n745) L3-rep1), GSM1534087(lin-35[JA1507(n745) L3-rep2)*.

## ACKNOWLEDGEMENTS

This work was supported by the NIH R35GM119775 to T.A.M and GM044619 to Gary Ruvkun

## TABLES, FIGURES AND SUPPORTING INFORMATION

**S1 Table:** Top 50 upregulated Genes in *lin-15B(n744*), mutant in starved L1 animals. The 50 most highly upregulated genes in *lin-15(n744)* mutants during the L1 stage. The log2 Fold-change and the corresponding false discovery rate (FDR), value for each gene that is listed is shown. The gene expression data was derived from our analysis of mRNA seq data sets *[GSM4697084-N2 - Rep 1/L1, GSM4697085-N2 - Rep 2/L1 against GSM4697102-lin-15b(n744)-rep1/L1, GSM4697105-lin-15b(n744)-rep2/L1*], that were obtained from the NCBI Geo collection.

| Gene | log2 fold change | FDR |
| --- | --- | --- |
| *pals-5* | 9.06 | 5.80E-07 |
| *pals-11* | 8.66 | 1.51E-05 |
| *pals-12* | 8.01 | 1.01E-06 |
| *F15H10.9* | 7.84 | 0.00602431 |
| *pals-10* | 7.78 | 0.0035545 |
| *Y105C5A.1269* | 7.63 | 3.18E-06 |
| *H05C05.3* | 7.54 | 4.39E-05 |
| *F15H10.12* | 7.16 | 0.00781933 |
| *Y82E9BL.19* | 7.06 | 2.88E-05 |
| *pals-29* | 6.99 | 1.63E-05 |
| *fbxa-75* | 6.97 | 2.01E-05 |
| *F15H10.10* | 6.72 | 0.00014014 |
| *F15H10.6* | 6.59 | 9.49E-05 |
| *C08E3.15* | 6.44 | 0.01574134 |
| *pes-2.1* | 6.43 | 0.28750516 |
| *pals-9* | 6.36 | 7.43E-05 |
| *Y54H5A.5* | 6.25 | 8.87E-09 |
| *F57G4.11* | 6.18 | 2.06E-05 |
| *Y45F10C.6* | 6.10 | 2.64E-05 |
| *Y61B8B.2* | 6.05 | 0.00502144 |
| *ZC434.8* | 5.84 | 3.27E-09 |
| *math-1* | 5.55 | 0.00161985 |
| *srw-85* | 5.51 | 1.43E-07 |
| *cpg-2* | 5.44 | 6.18E-07 |
| *Y51H7C.12* | 5.37 | 0.03414094 |
| *F37D6.3* | 5.37 | 4.15E-08 |
| *sdz-6* | 5.26 | 0.05818951 |
| *Y38E10A.3* | 5.26 | 0.19355176 |
| *F40G12.11* | 5.25 | 0.17805195 |
| *ZK250.14* | 5.25 | 0.00148968 |
| *Y57G7A.2* | 5.24 | 0.20756677 |
| *skr-9* | 5.19 | 0.11960343 |
| *sri-68* | 5.15 | 0.00162513 |
| *C08F11.7* | 5.14 | 0.00026807 |
| *F22E5.20* | 5.14 | 0.31126046 |
| *pals-3* | 5.11 | 0.00014288 |
| *Y39H10B.3* | 5.07 | 1.77E-06 |
| *F20E11.17* | 5.04 | 0.00089112 |
| *math-5* | 5.03 | 0.00911578 |
| *pals-28* | 4.99 | 0.00452157 |
| *fbxb-90* | 4.97 | 0.27388713 |
| *ZK666.11* | 4.94 | 0.03078516 |
| *C29A12.1* | 4.92 | 9.83E-08 |
| *T05A8.2* | 4.78 | 0.01574134 |
| *E02H9.3* | 4.75 | 9.83E-08 |
| *ucr-2.3* | 4.74 | 8.87E-09 |
| *C38D4.1* | 4.72 | 5.09E-07 |
| *F40E12.2* | 4.71 | 0.00044008 |
| *srg-53* | 4.70 | 0.0021712 |
| *pals-4* | 4.67 | 0.00050884 |
| *pals-14* | 4.67 | 2.95E-06 |

**S2 Table:**
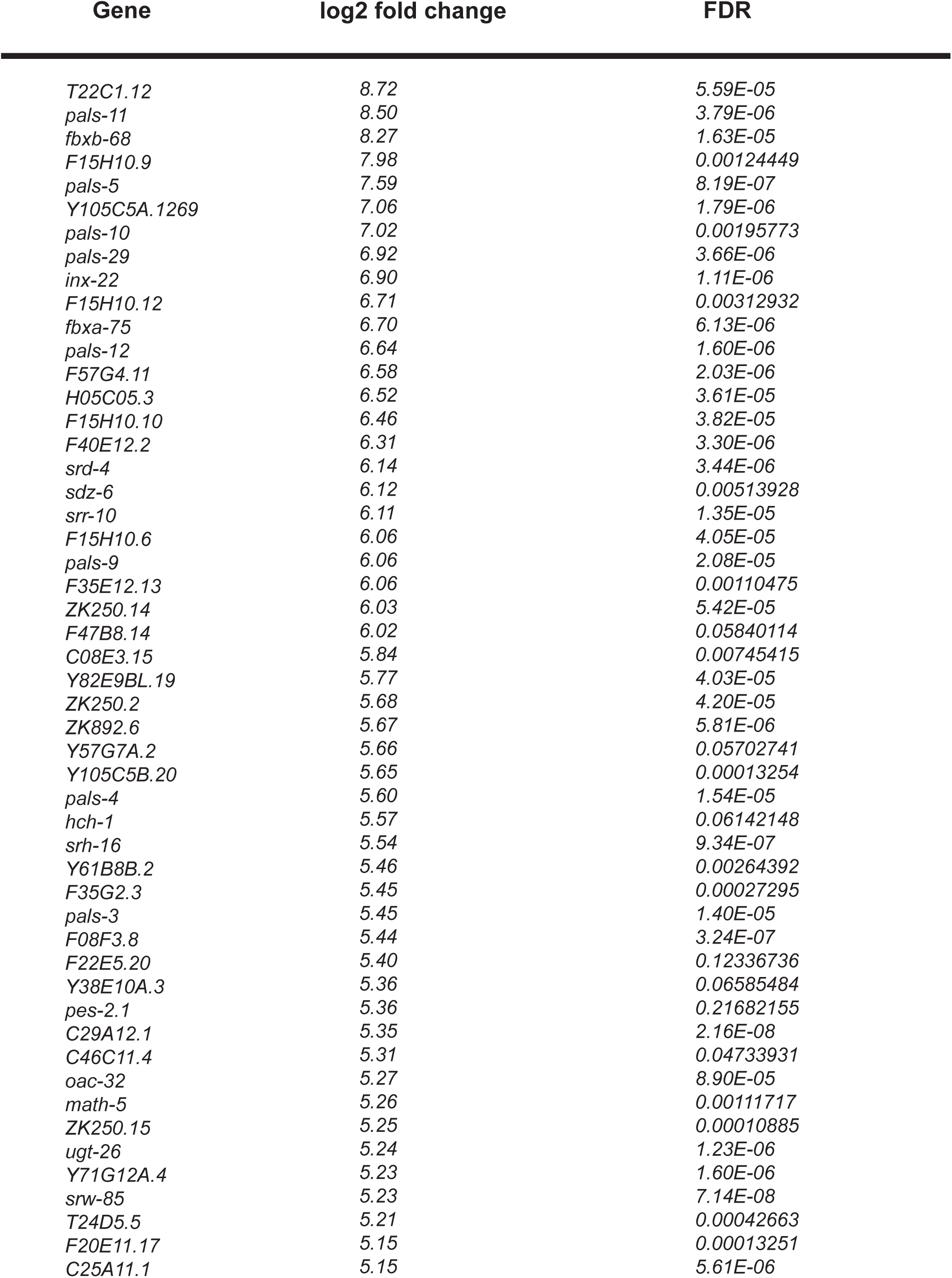
Top 50 upregulated Genes in *lin-35(n745*), mutant in starved L1 animals. The 50 most highly upregulated genes in *lin-35(n745)* mutants during the L1 stage. The log2 Fold-change and the corresponding false discovery rate (FDR), value for each gene that is listed is shown. The gene expression data was derived from our analysis of mRNA seq data sets *GSM4697089-lin-35[JA1507(n745) rep1/L1, GSM4697090-lin-35[JA1507(n745) rep1/L1],* that were obtained from the NCBI Geo collection.

| Gene | log2 fold change | FDR |
| --- | --- | --- |
| T22C1.12 | 8.72 | 5.59E-05 |
| pals-11 | 8.50 | 3.79E-06 |
| fbxb-68 | 8.27 | 1.63E-05 |
| F15H10.9 | 7.98 | 0.00124449 |
| pals-5 | 7.59 | 8.19E-07 |
| Y105C5A.1269 | 7.06 | 1.79E-06 |
| pals-10 | 7.02 | 0.00195773 |
| pals-29 | 6.92 | 3.66E-06 |
| inx-22 | 6.90 | 1.11E-06 |
| F15H10.12 | 6.71 | 0.00312932 |
| fbxa-75 | 6.70 | 6.13E-06 |
| pals-12 | 6.64 | 1.60E-06 |
| F57G4.11 | 6.58 | 2.03E-06 |
| H05C05.3 | 6.52 | 3.61E-05 |
| F15H10.10 | 6.46 | 3.82E-05 |
| F40E12.2 | 6.31 | 3.30E-06 |
| srd-4 | 6.14 | 3.44E-06 |
| sdz-6 | 6.12 | 0.00513928 |
| srr-10 | 6.11 | 1.35E-05 |
| F15H10.6 | 6.06 | 4.05E-05 |
| pals-9 | 6.06 | 2.08E-05 |
| F35E12.13 | 6.06 | 0.00110475 |
| ZK250.14 | 6.03 | 5.42E-05 |
| F47B8.14 | 6.02 | 0.05840114 |
| C08E3.15 | 5.84 | 0.00745415 |
| Y82E9BL.19 | 5.77 | 4.03E-05 |
| ZK250.2 | 5.68 | 4.20E-05 |
| ZK892.6 | 5.67 | 5.81E-06 |
| Y57G7A.2 | 5.66 | 0.05702741 |
| Y105C5B.20 | 5.65 | 0.00013254 |
| pals-4 | 5.60 | 1.54E-05 |
| hch-1 | 5.57 | 0.06142148 |
| srh-16 | 5.54 | 9.34E-07 |
| Y61B8B.2 | 5.46 | 0.00264392 |
| F35G2.3 | 5.45 | 0.00027295 |
| pals-3 | 5.45 | 1.40E-05 |
| F08F3.8 | 5.44 | 3.24E-07 |
| F22E5.20 | 5.40 | 0.12336736 |
| Y38E10A.3 | 5.36 | 0.06585484 |
| pes-2.1 | 5.36 | 0.21682155 |
| C29A12.1 | 5.35 | 2.16E-08 |
| C46C11.4 | 5.31 | 0.04733931 |
| oac-32 | 5.27 | 8.90E-05 |
| math-5 | 5.26 | 0.00111717 |
| ZK250.15 | 5.25 | 0.00010885 |
| ugt-26 | 5.24 | 1.23E-06 |
| Y71G12A.4 | 5.23 | 1.60E-06 |
| srw-85 | 5.23 | 7.14E-08 |
| T24D5.5 | 5.21 | 0.00042663 |
| F20E11.17 | 5.15 | 0.00013251 |
| C25A11.1 | 5.15 | 5.61E-06 |

**S3 Table:** Top 50 upregulated Genes in *lin-35(n745*), mutant at the L3 Stage. The top 50 most highly upregulated genes in *lin-35(n745)* mutants during the L3 stage. The log2 Fold-change and the corresponding false discovery rate (FDR), value for each gene that is listed is shown. [*GSM1534084-N2-rep1 L3, GSM1534085-N2-rep2 compared against GSM1534086-lin-35[JA1507(n745) rep1 L3 and GSM1534087-lin-35[JA1507(n745) rep1 L3],* that were obtained from the NCBI Geo collection.

| Gene | log2 fold change | FDR |
| --- | --- | --- |
| *fbxa-75* | 11.55 | 4.28E-04 |
| *pals-3* | 11.46 | 3.47E-04 |
| *pals-8* | 10.93 | 6.37E-04 |
| *pals-11* | 9.69 | 1.15E-03 |
| *F26F2.4* | 9.47 | 5.38E-03 |
| *F15H10.9* | 9.32 | 1.44E-02 |
| *pals-29* | 9.10 | 4.99E-04 |
| *oac-56* | 8.82 | 2.03E-02 |
| *B0507.15* | 8.81 | 4.53E-03 |
| *F40E12.2* | 8.79 | 5.73E-04 |
| *sdz-6* | 8.72 | 1.27E-02 |
| *fbxa-165* | 8.68 | 2.21E-04 |
| *F26F2.5* | 8.59 | 2.10E-03 |
| *F15H10.5* | 8.51 | 3.41E-02 |
| *srd-4* | 8.38 | 4.62E-04 |
| *E03H4.9* | 8.33 | 2.15E-03 |
| *Y54G2A.13* | 8.32 | 6.55E-03 |
| *srh-16* | 8.30 | 1.20E-04 |
| *K06A5.2* | 8.14 | 4.63E-02 |
| *F57G4.1* | 8.11 | 1.01E-02 |
| *C43D7.4* | 7.96 | 1.89E-02 |
| *F57G4.11* | 7.89 | 1.63E-04 |
| *pals-5* | 7.89 | 3.47E-04 |
| *Y39H10B.3* | 7.84 | 7.04E-05 |
| *fbxa-7* | 7.77 | 1.17E-03 |
| *math-16* | 7.77 | 7.70E-02 |
| *C04B4.2* | 7.75 | 1.04E-01 |
| *F45E4.6* | 7.74 | 8.46E-03 |
| *pals-9* | 7.72 | 9.99E-04 |
| *T27E7.9* | 7.67 | 3.72E-02 |
| *T24A6.7* | 7.63 | 2.10E-02 |
| *pals-7* | 7.51 | 2.97E-03 |
| *F37D6.3* | 7.48 | 9.45E-06 |
| *H04D03.6* | 7.43 | 3.21E-02 |
| *B0281.8* | 7.43 | 2.53E-02 |
| *pals-2* | 7.38 | 7.04E-05 |
| *F55G11.3* | 7.34 | 4.63E-02 |
| *math-17* | 7.34 | 4.79E-02 |
| *F15H10.6* | 7.33 | 1.45E-03 |
| *F26F2.3* | 7.13 | 2.69E-05 |
| *fbxb-68* | 7.10 | 2.79E-02 |
| *F36H5.10* | 7.09 | 1.55E-02 |
| *F35G2.3* | 7.06 | 1.01E-02 |
| *srg-31* | 7.04 | 2.75E-03 |
| *sri-68* | 7.03 | 3.12E-02 |
| *pals-33* | 6.97 | 9.45E-06 |
| *pals-4* | 6.89 | 1.15E-03 |
| *fbxa-173* | 6.85 | 1.42E-01 |
| *Y61B8B.2* | 6.82 | 1.83E-02 |
| *F20E11.17* | 6.79 | 2.83E-03 |

**S1 Fig.**
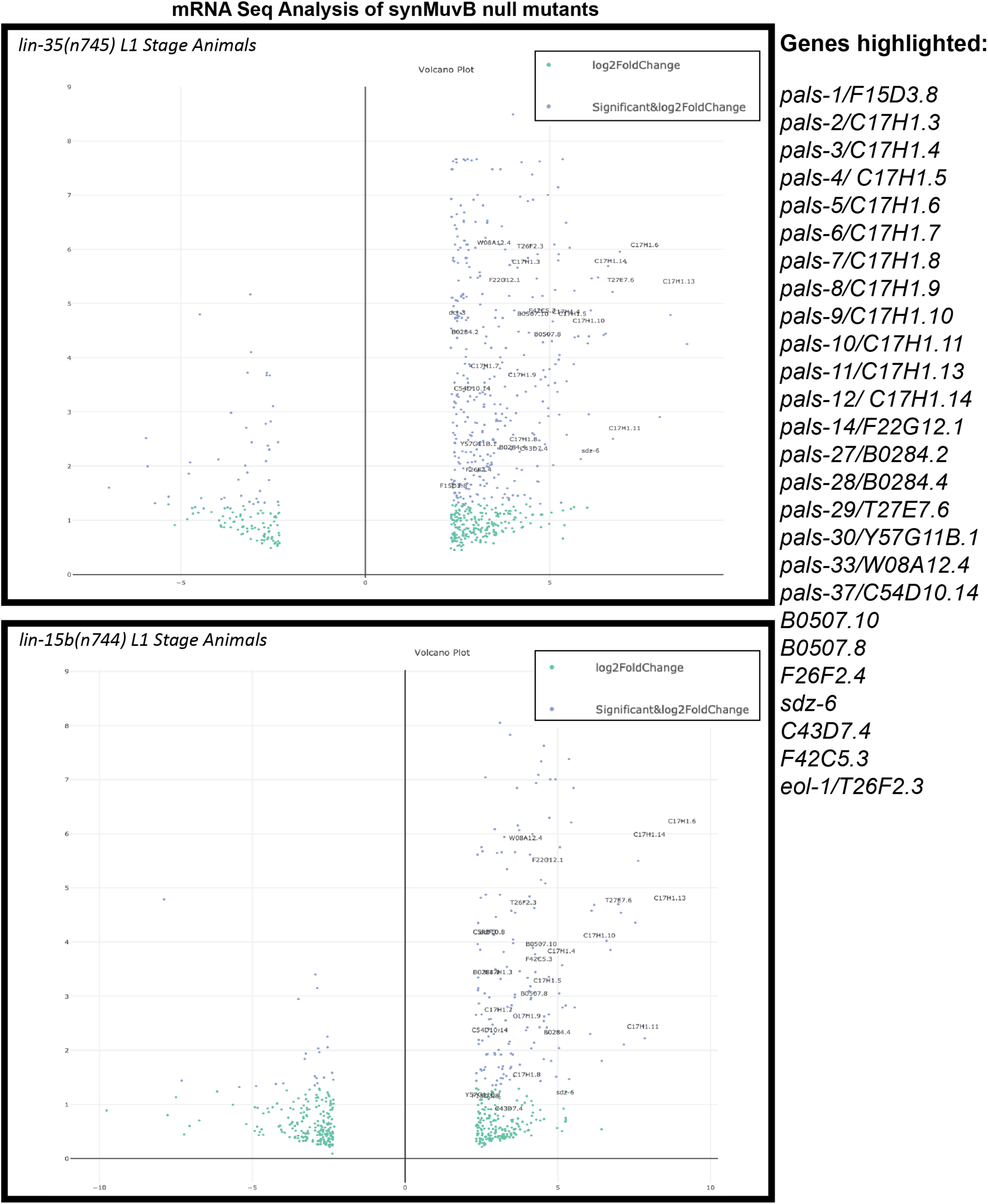
Gene expression analysis reveals antiviral defense genes are upregulated in SynMuv B mutants. Volcano plots of a gene expression analysis done on mRNA seq data available through NCBI Geo for the null mutants of *lin-35(n745); glp-4(bn2)* and *glp-4(bn2); lin-15b(n744).* Highlighted are those GeneIDs belonging to genes that have been previously reported to play a role in antiviral defense response in worms. These genes are also listed on the right. Each of the highlighted GeneIDs have a cutoff of a logFC of 2.32 (i.e. a 5-fold change).

**S2 Fig.**
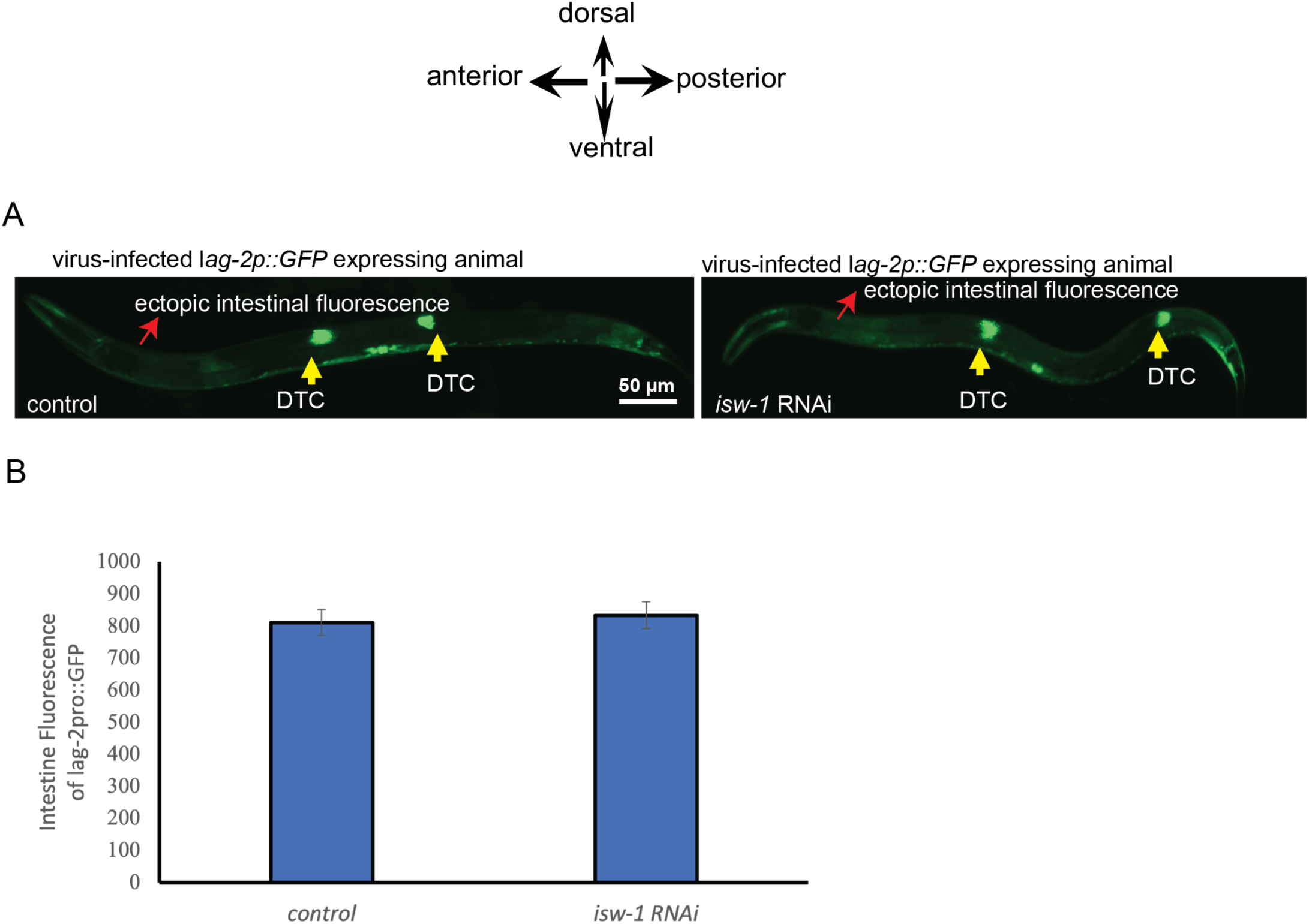
Ectopic *lag-2p::GFP* expression in the intestinal cells upon virus-infection remains unaltered upon removal of *isw-1* activity by RNAi. (A). Fluorescent micrographs showing two *lag-2p::GFP* expressing animals that are infected with the Orsay virus raised on *E. coli* that express double stranded RNA against either empty vector (control) or the synMuv suppressor gene *isw-1*. Expression of *lag-2p::GFP* within the distal tip cells is indicated by yellow arrow heads and the ectopic fluorescence of *lag-2p::GFP* in the intestine is also indicated. (B). Quantification of the fluorescence intensity of *lag-2p::GFP* within the intestine of animals that are infected with the Orsay virus raised on *E. coli* that express double stranded RNA against either the L4440 empty vector (labelled as control) or *isw-1* is shown. A schematic of the general orientation of the worm is shown above the image panels. Scale bar is indicated.

**S3 Fig.**
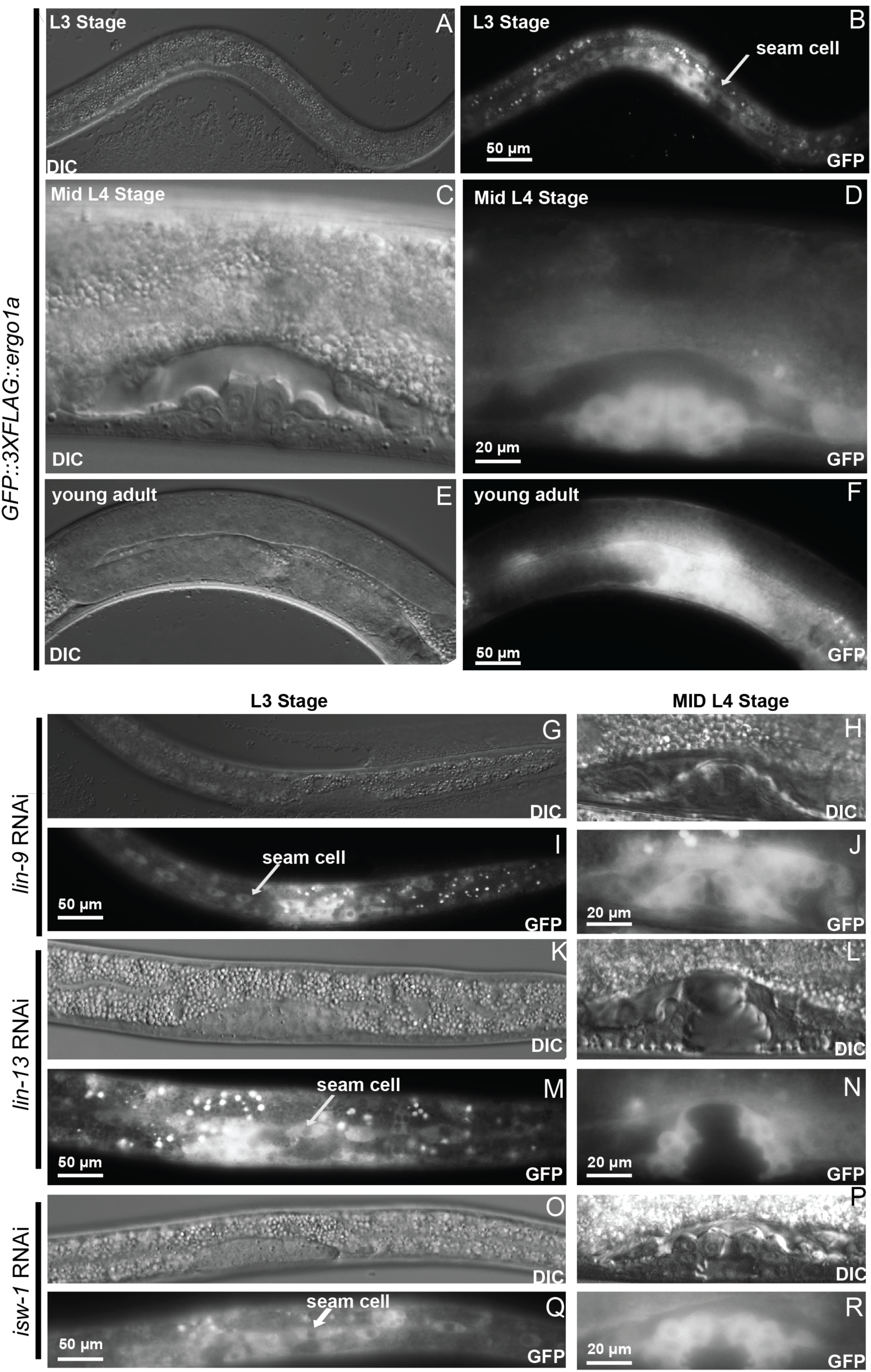
Spatiotemporal expression pattern of ERGO-1 remains unperturbed by the loss of function of synMuv B genes. (A-F). Brightfield and GFP channel micrographs depicting the temporal expression pattern of *ergo-1::GFP* from a transgenic strain that expresses a full length ERGO-1 PIWI protein fused at its C terminus to GFP. (G-R). *ergo-1::GFP* expression after synMuv B (*lin-9 or lin-13*), or Muv suppressor (*isw-1*) RNAi. Developmental stage is labelled. Scale bar is indicated.

**S4 Fig.**
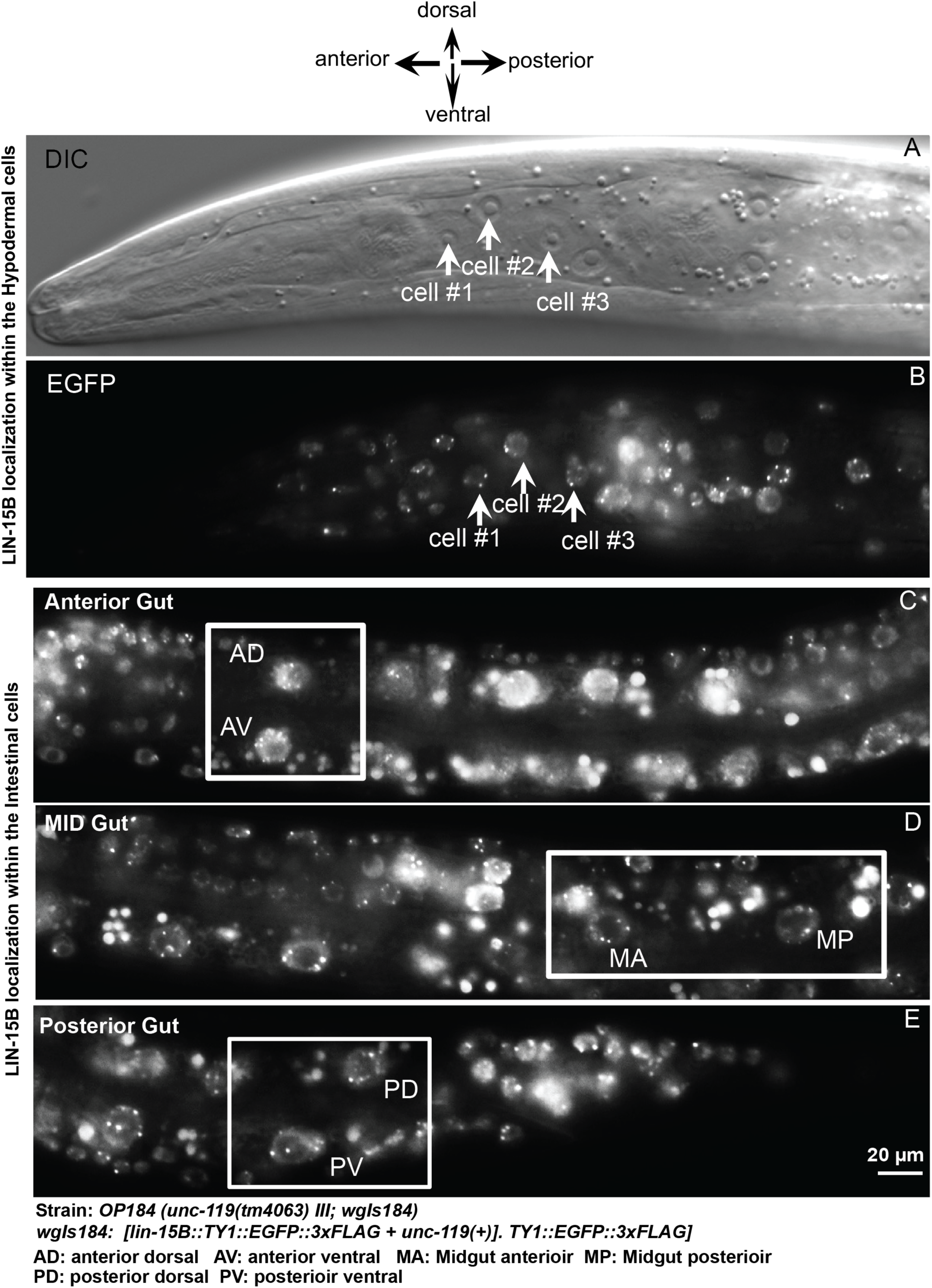
LIN-15B C-terminal protein fusion::EGFP localization in hypodermal and intestinal cells in *C. elegans*. (A-E). Brightfield and EGFP micrographs of LIN-15B localization in representative hypodermal (panels A & B), intestinal nuclei (panels C,D &E). Abbreviations for the different intestinal cells that are boxed is provided below the image panels. Scale bar is indicated.

**S5 Fig.**
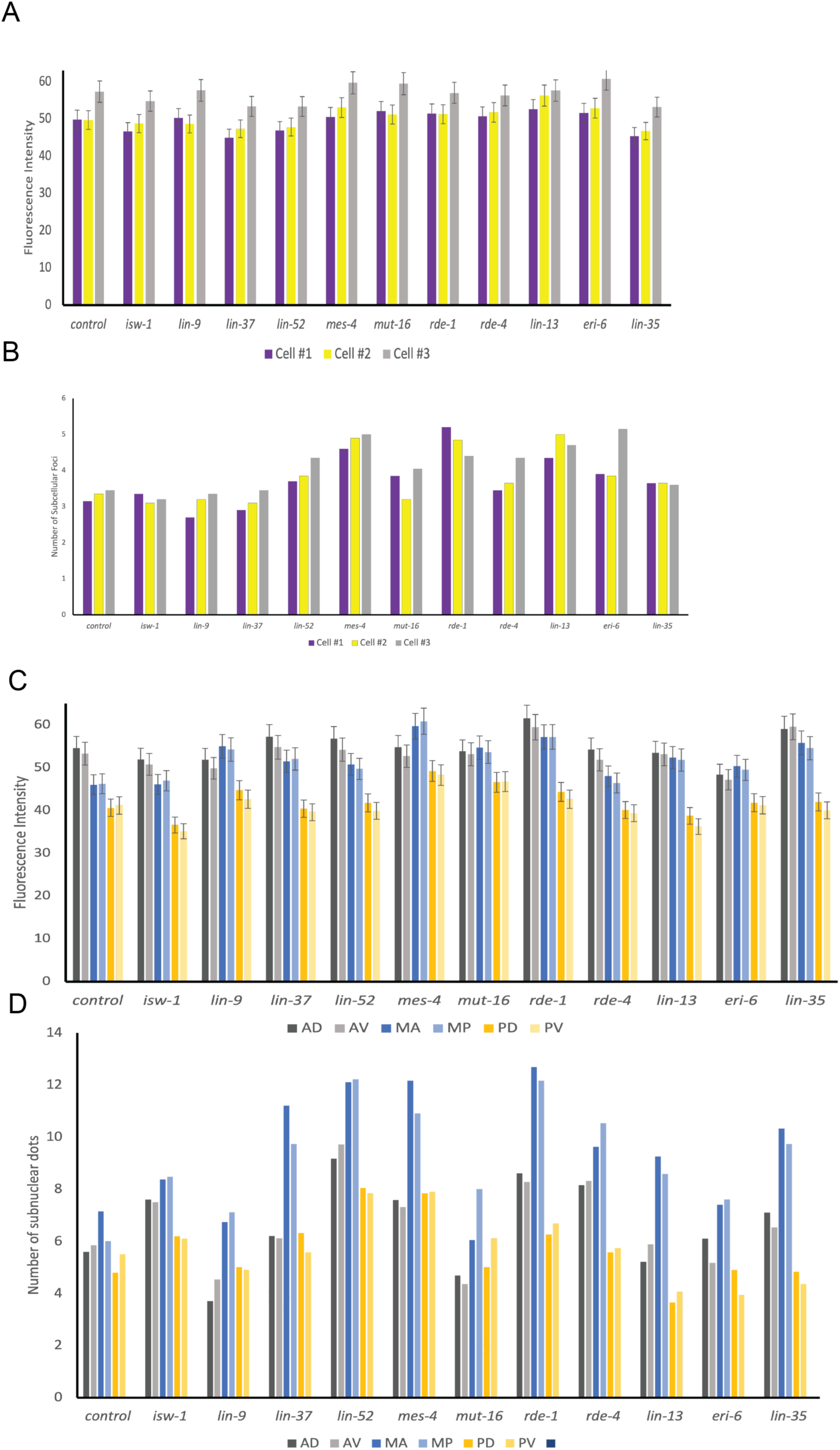
Quantitation of LIN-15B::EGFP fluorescence intensity and number of subnuclear foci. (A&B). Quantification of EGFP fluorescence and number of subnuclear LIN-15B foci in 3 representative hypodermal cells for synMuv B (*lin-9, lin-13, lin-35, lin-37 and lin-52*), Eri associated-gene (*eri-6*), synMuv suppressors (*mes-4, isw-1*), RNAi defective genes (*rde-1, rde-4, mut-16*) under RNAi conditions. (C&D). Quantification of EGFP fluorescence and number of subnuclear LIN-15B foci within 6 representative intestinal cells for synMuv B (*lin-9, lin-13, lin-35, lin-37 and lin-52*), Eri associated-gene (*eri-6*), synMuv suppressors (*mes-4, isw-1*), RNAi defective genes (*rde-1, rde-4, mut-16*) under RNAi conditions. In the graphs shown in C and D the abbreviations AD refers to anterior dorsal cell, AV refers to anterior ventral cell, MA refers to midgut anterior cell and MP refers to midgut posterior cell, PD refers to posterior gut dorsal cell and PV refers to posterior gut ventral cell.

**S6 Fig.**
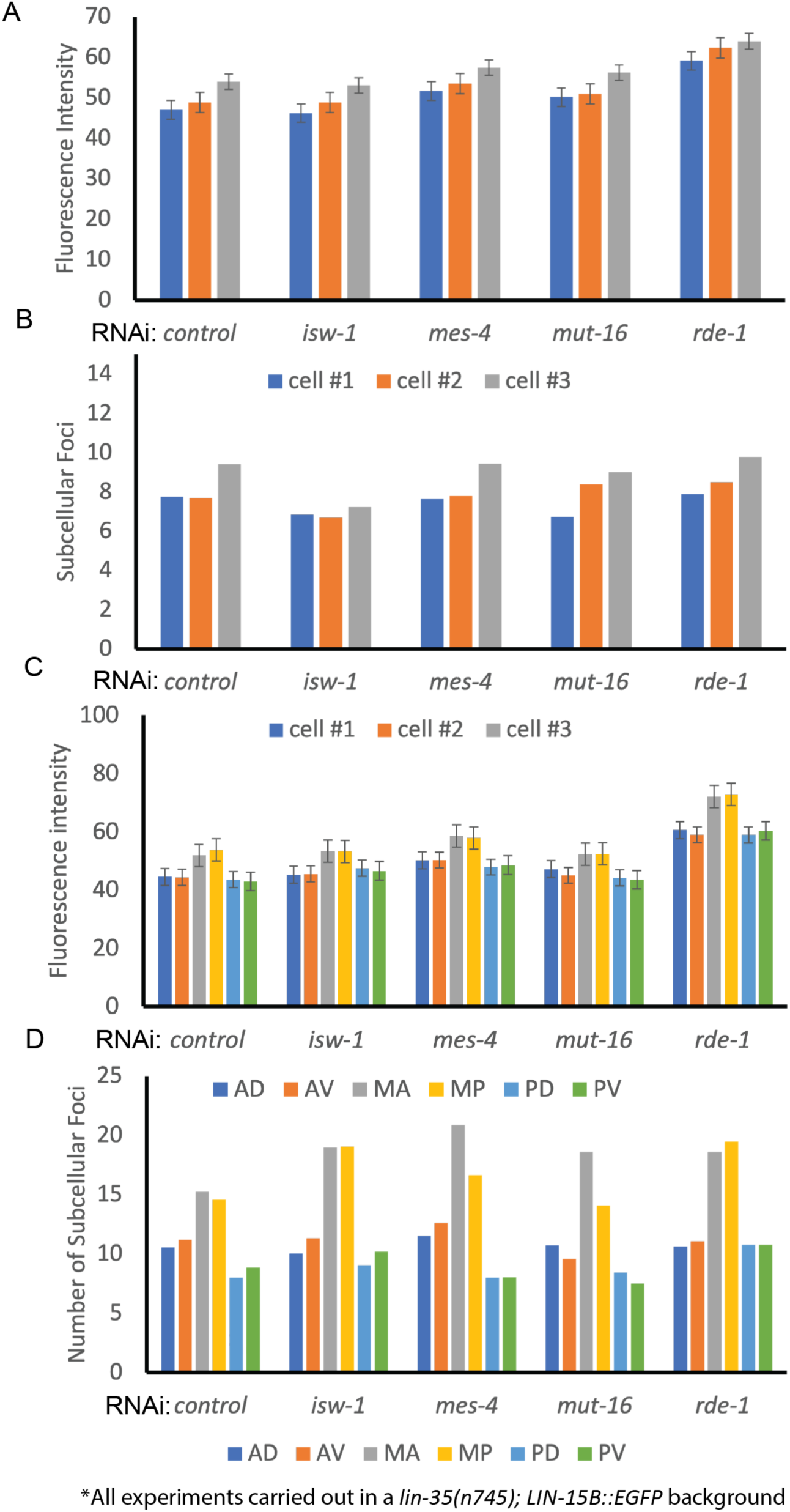
Quantification of LIN-15B::EGFP fusion protein fluorescence intensity and number of subnuclear foci in *lin-35(n745)* null mutants. (A&B). EGFP fluorescence and number of subnuclear LIN-15B foci within 3 representative hypodermal cells under RNAi conditions targeting L4440 vector (control), RNAi defective genes (*rde-1, mut-16*) and Muv suppressor (*isw-1 and mes-4*). (C&D). Intensity of EGFP fluorescence and number of subnuclear LIN-15B foci within 6 representative intestinal cells where the abbreviations AD refers to anterior dorsal cell, AV refers to anterior ventral cell, MA refers to midgut anterior cell and MP refers to midgut posterior cell, PD refers to posterior gut dorsal cell and PV refers to posterior gut ventral cell. RNAi was performed targeting L4440 vector (control), RNAi defective genes (*rde-1, mut-16*) and Muv suppressor genes (*isw-1 and mes-4*). These experiments were carried out in a *lin-35(n745)* null background.

**S7 Fig.**
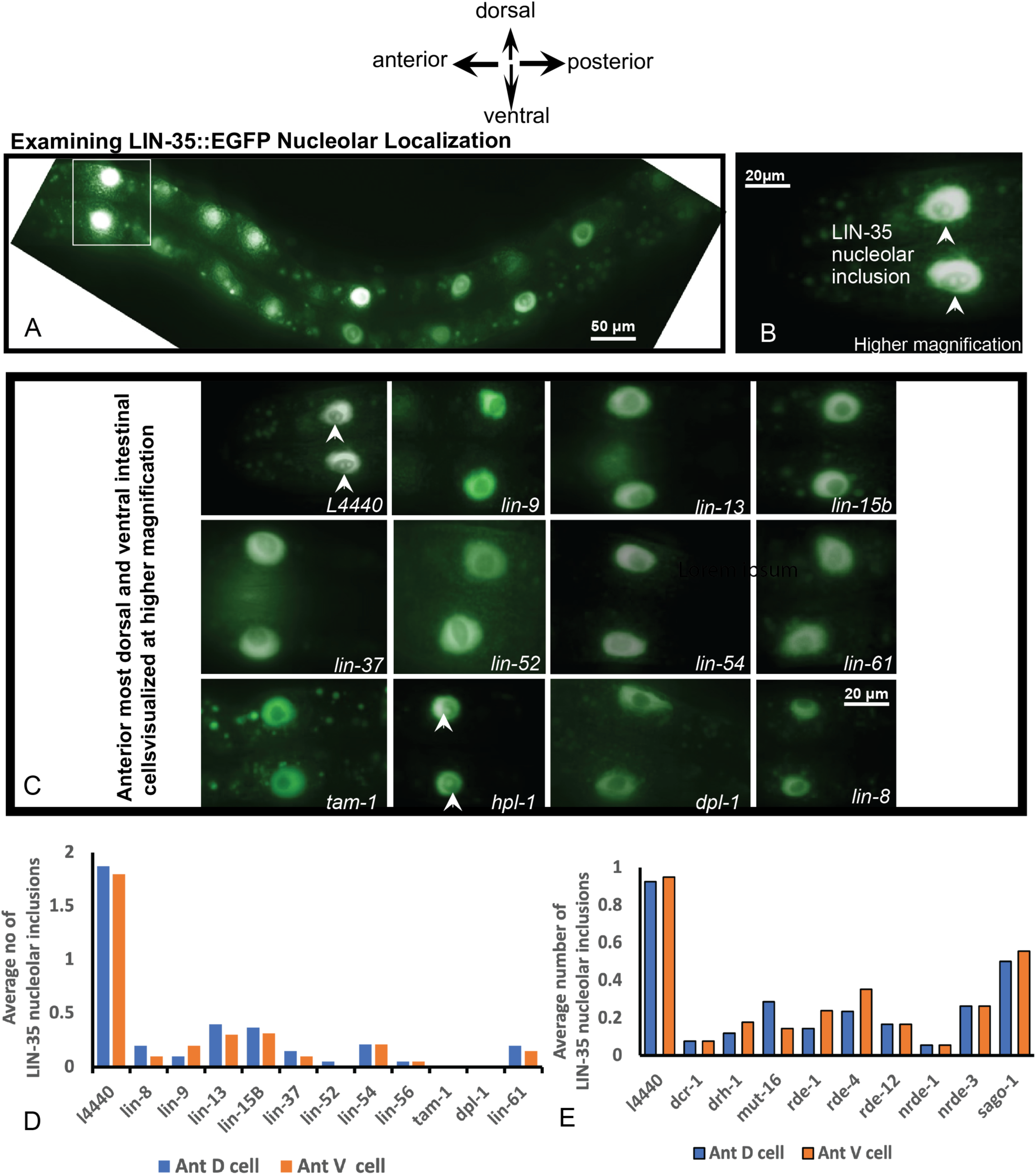
LIN-35::EGFP localization in the nucleolus of *C. elegans* intestinal cells. (A & B). Fluorescent micrographs of LIN-35::EGFP fusion protein localization in the intestinal nucleoli of wild-type animals. Magnification used to visualize the cells is indicated. The image shown in panel B are the same cells that are boxed in panel A, visualized under a higher magnification. (C). Fluorescent micrographs of LIN-35::EGFP localization within the intestinal cells (boxed in panel A), of the F1 progeny of worms raised on *E. coli* expressing double stranded RNA against either empty vector (l4440) or various synMuv B genes (*lin-9, lin-13, lin-15b, lin-37, lin-52, lin-54, lin-61, tam-1, hpl-1 and dpl-1*) or a synMuv A gene (*lin-8*) is shown. Arrow heads indicate the presence of a nucleolar inclusion of LIN-35::EGFP. (D). Quantification of the number of LIN-35::EGFP nucleolar inclusions that are seen in animals raised on *E. coli* expressing double stranded RNA against either empty vector (l4440) or various synMuv B genes (*lin-9, lin-13, lin-15b, lin-37, lin-52, lin-54, lin-61, tam-1, hpl-1 and dpl-1*) or a synMuv A gene (*lin-8*) is shown. (E). Quantification of the number of LIN-35::EGFP nucleolar inclusions that are seen in animals raised on *E. coli* expressing double stranded RNA against either empty vector (l4440) or various genes that are critical for the worms ability to perform RNAi. Scale bar is indicated.

**S8 Fig.**
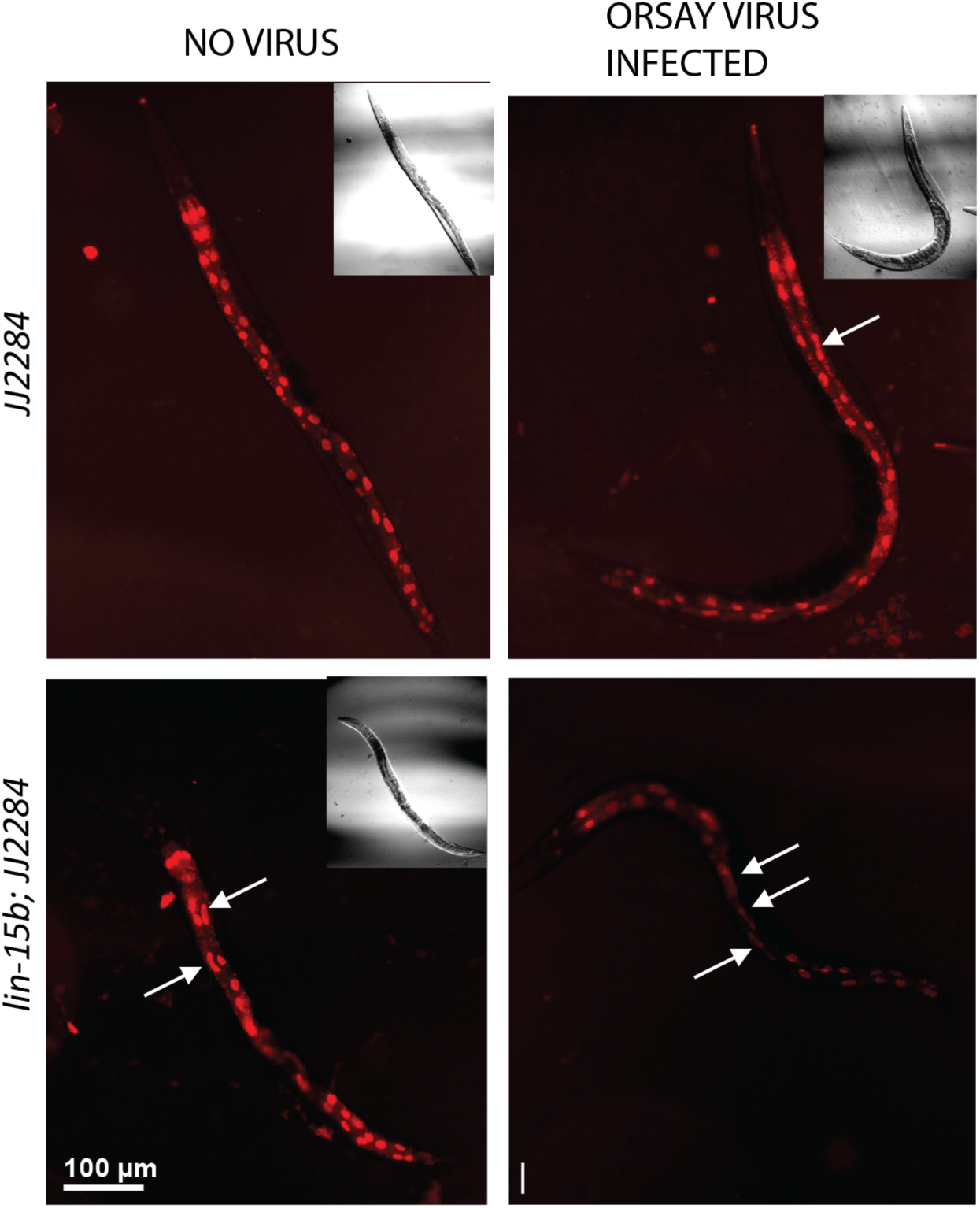
*lin-15b(-)* animals exhibit altered intestinal nuclear morphology. Top panel shows wild-type *JJ2284* animals under no virus and Orsay virus-infected conditions. Lower panel shows a second independent line of *lin-15b(W485*)*; JJ2284 animals under no virus and Orsay virus-infected conditions. White arrows indicate elongated intestinal nuclei that are elongated. Scale bar is indicated.

**S9 Fig.**
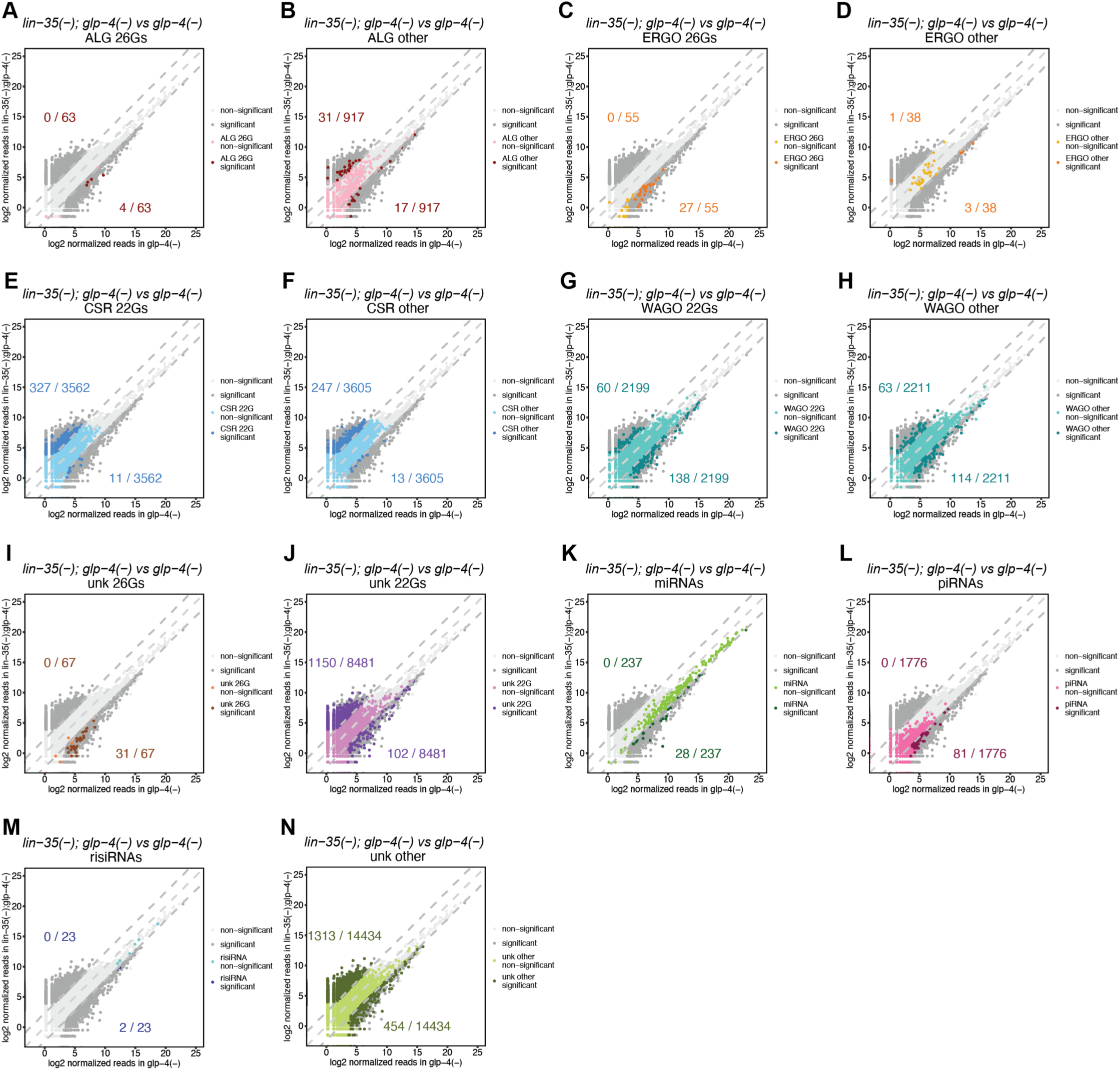
Comparisons of normalized reads of indicated classes of small RNAs in *lin-35(n745); glp-4(bn2)* with *glp-4(bn2)*. Dark grey data points represent small RNAs that are differentially expressed by 5-fold and adjusted p value < 0.05. Colored data points represent indicated classes of small RNAs. The rest of the small RNAs are represented as light grey data points. Lines denoting equal, 5-fold increased and 5-fold decreased expression are shown as light grey dashed lines. The numbers indicate the number of small RNAs upregulated in *lin-35(n745); glp-4(bn2)* (upper) and *glp-4(bn2)* (lower).

**S10 Fig.**
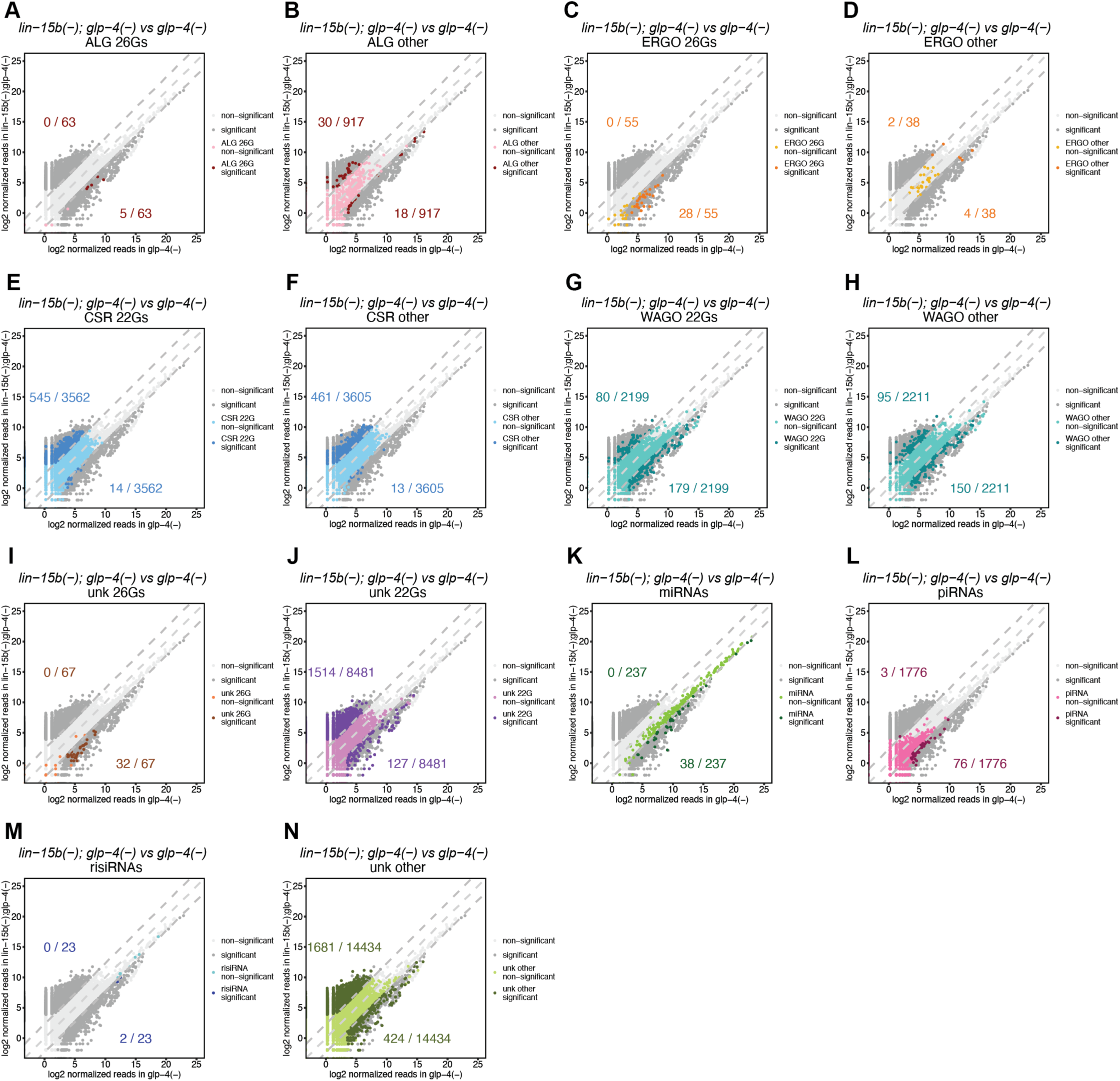
Comparisons of normalized reads of indicated classes of small RNAs in *lin-15b(n744); glp-4(bn2)* with *glp-4(bn2)*. Dark grey data points represent small RNAs that are differentially expressed by 5-fold and adjusted p value < 0.05. Colored data points represent indicated classes of small RNAs. The rest of the small RNAs are represented as light grey data points. Lines denoting equal, 5-fold increased and 5-fold decreased expression are shown as light grey dashed lines. The numbers indicate the number of small RNAs upregulated in *lin-15b(n744); glp-4(bn2)* (upper) and *glp-4(bn2)* (lower).

**S11 Fig.**
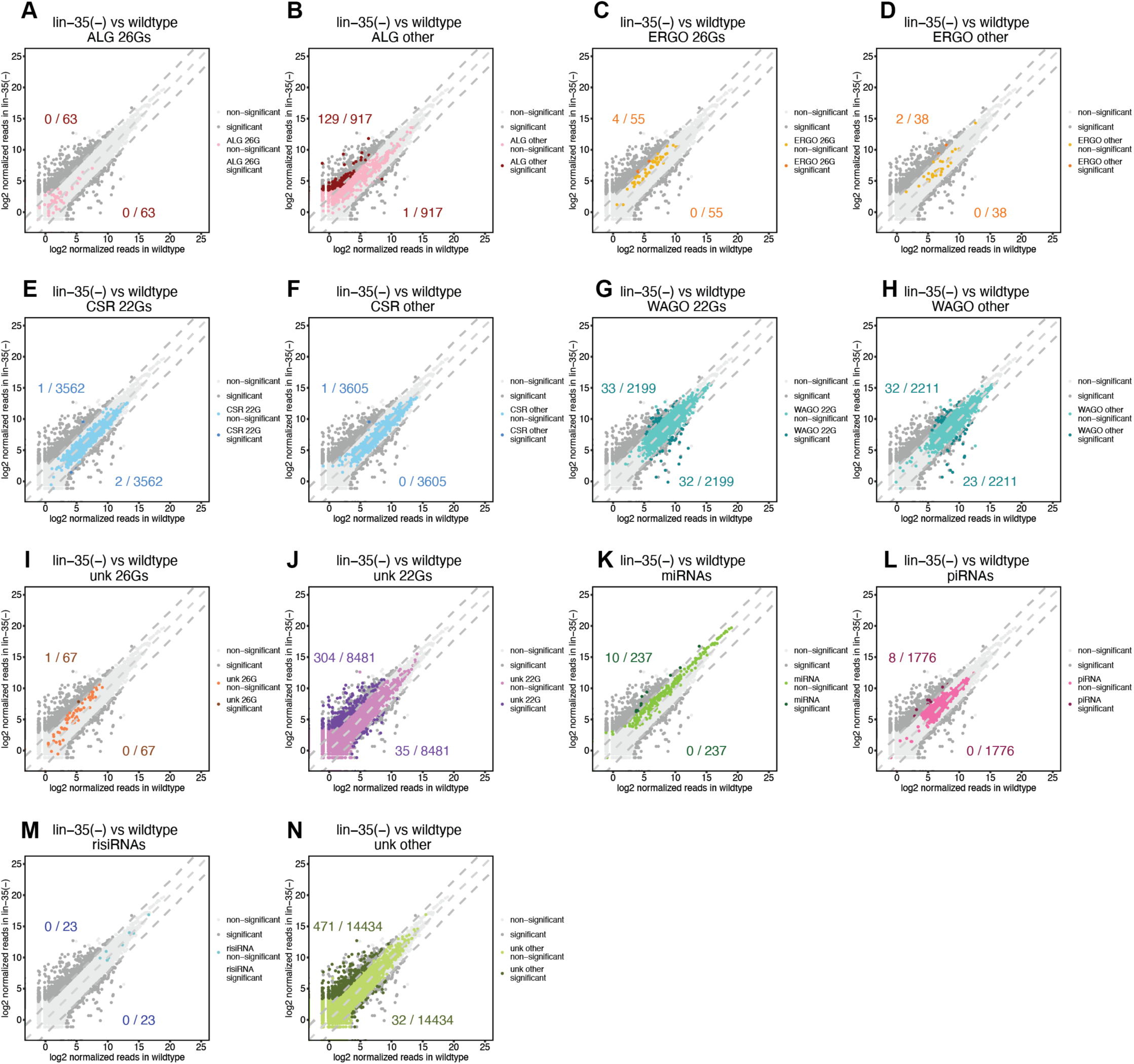
Comparisons of normalized reads of indicated classes of small RNAs in *lin-35(n745)* with wildtype. Dark grey data points represent small RNAs that are differentially expressed by 5-fold and adjusted p value < 0.05. Colored data points represent indicated classes of small RNAs. The rest of the small RNAs are represented as light grey data points. Lines denoting equal, 5-fold increased and 5-fold decreased expression are shown as light grey dashed lines. The numbers indicate the number of small RNAs upregulated in *lin-35(n745)* (upper) and wildtype (lower).

**S12 Fig.**
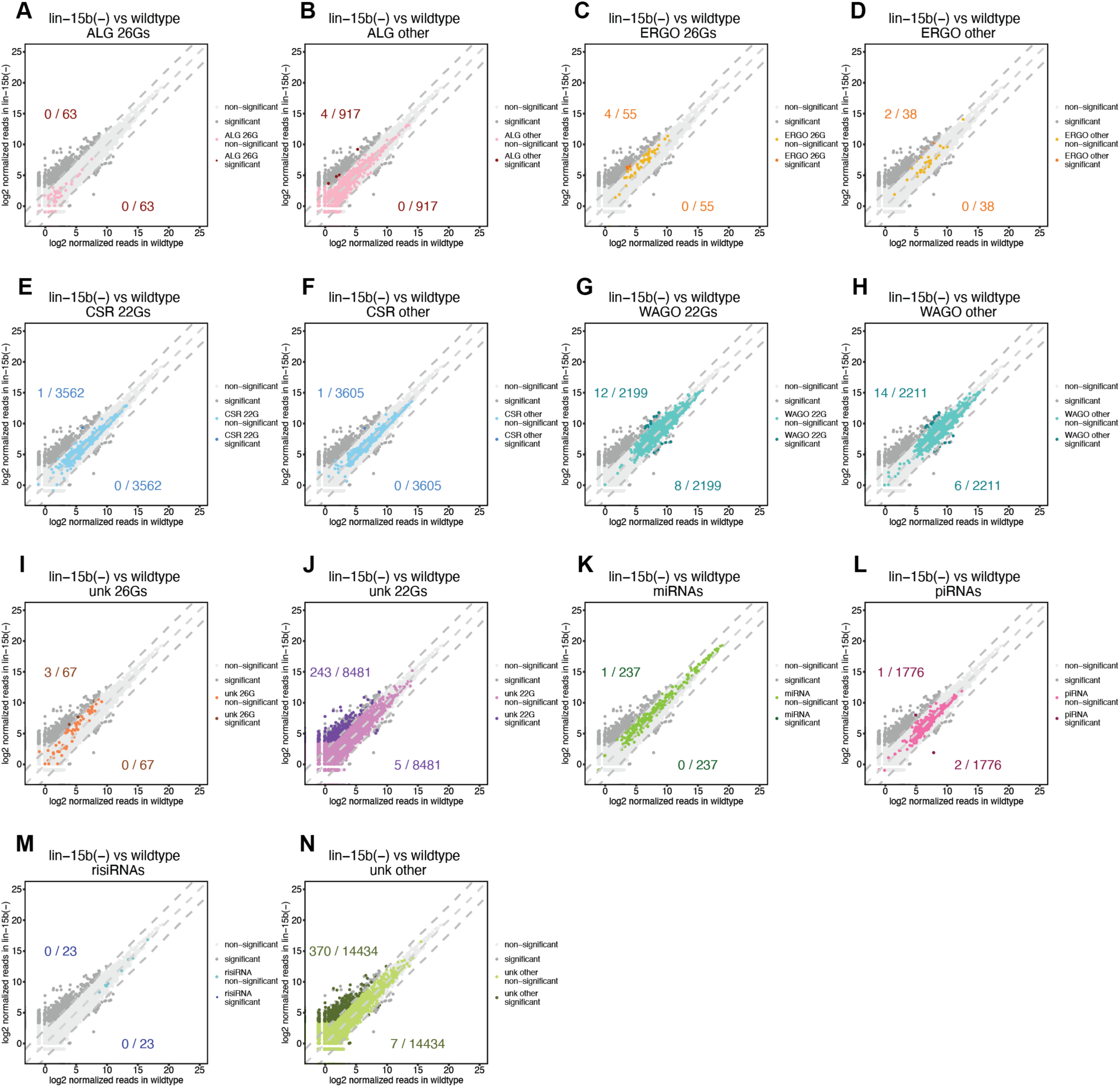
Comparisons of normalized reads of indicated classes of small RNAs in *lin-15b(n744)* with wildtype. Dark grey data points represent small RNAs that are differentially expressed by 5-fold and adjusted p value < 0.05. Colored data points represent indicated classes of small RNAs. The rest of the small RNAs are represented as light grey data points. Lines denoting equal, 5-fold increased and 5-fold decreased expression are shown as light grey dashed lines. The numbers indicate the number of small RNAs upregulated in *lin-15b(n744)*(upper) and wildtype (lower).

**S13 Fig.**
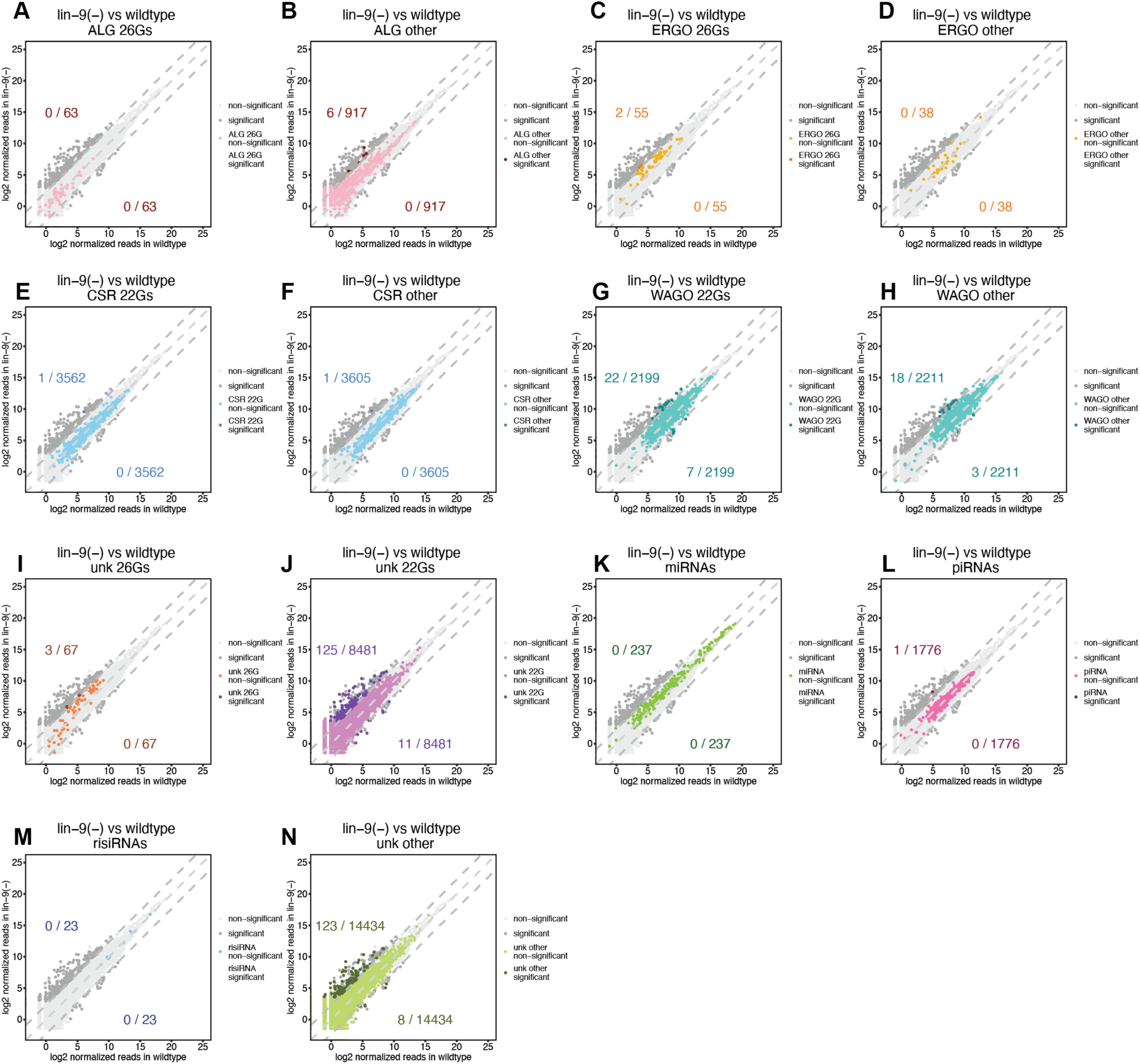
Comparisons of normalized reads of indicated classes of small RNAs in *lin-9(n112)* with wildtype. B Dark grey data points represent small RNAs that are differentially expressed by 5-fold and adjusted p value < 0.05. Colored data points represent indicated classes of small RNAs. The rest of the small RNAs are represented as light grey data points. Lines denoting equal, 5-fold increased and 5-fold decreased expression are shown as light grey dashed lines. The numbers indicate the number of small RNAs upregulated in *lin-9(n112)* (upper) and wildtype (lower).

**S14 Fig.**
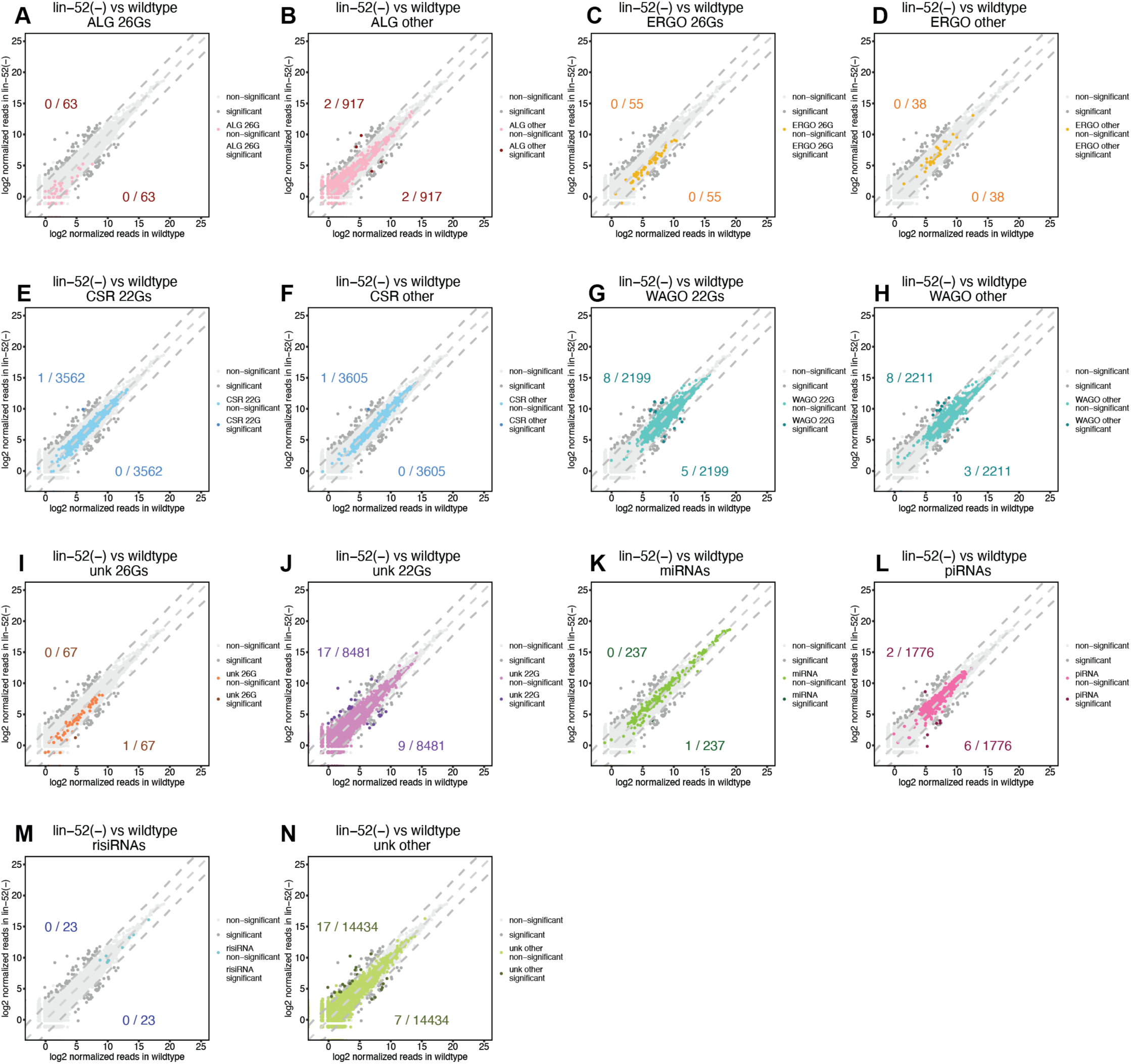
Comparisons of normalized reads of indicated classes of small RNAs in *lin-52(n771)* with wildtype. Dark grey data points represent small RNAs that are differentially expressed by 5-fold and adjusted p value < 0.05. Colored data points represent indicated classes of small RNAs. The rest of the small RNAs are represented as light grey data points. Lines denoting equal, 5-fold increased and 5-fold decreased expression are shown as light grey dashed lines. The numbers indicate the number of small RNAs upregulated in *lin-52(n771)*(upper) and wildtype (lower).

## ADDITIONAL SUPPORTING INFORMATION DATA FILES: [uploaded separately]

**Additional supporting information data table 1. List of genes to which differentially regulated small RNAs map in *lin-35(n745); glp-4(bn2)* and *lin-15b(n744); glp-4(bn2)* in comparison with *glp-4(bn2).*** The siRNAs that are either Up or Down by a minimum factor of a 5-fold difference in the *lin-15(n744)* and *lin-35(n745)* mutant backgrounds are presented. See the labeled tabs for the genotype information.

**Additional supporting information data table 2. Gene expression changes in *lin-35(n745)* starved L1 animals.** Shown here are the list of significant gene expression changes via mRNA seq that are observed in the *lin-35(n745)* synMuv B mutant animals at the L1 stage. The data shown here was obtained from the NCBI geo collection, *GSM4697089 [lin-35(n745) starved L1-rep1)], GSM4697090 (lin-35(n745) starved L1-rep2)]*.

**Additional supporting information data table 3. Gene expression changes in *lin-35(n745)* starved L3-stage animals.** Shown here are the list of significant gene expression changes via mRNA seq that are observed in the *lin-35(n745)* synMuv B mutant animals at the L3-stage. The data shown here was obtained from the NCBI geo collection, *GSM1534086 (lin-35[JA1507(n745) L3-rep1), GSM1534087(lin-35[JA1507(n745) L3-rep2)]*.

**Additional supporting information data table 4. Gene expression changes in *lin-15(n744)* starved L1-stage animals.** Shown here are the list of significant gene expression changes via mRNA seq that are observed in the *lin-15(n744)* synMuv B mutant animals at the L1-stage. The data shown here was obtained from the NCBI geo collection, *GSM4697102 (lin-15B(n744) starved L1-rep1), GSM4697105 (lin-15B(n744) starved L1-rep2)]*.

